# 3D Bioprinting of Collagen-based Microfluidics for Engineering Fully-biologic Tissue Systems

**DOI:** 10.1101/2024.01.26.577422

**Authors:** Daniel J. Shiwarski, Andrew R. Hudson, Joshua W. Tashman, Ezgi Bakirci, Samuel Moss, Brian D. Coffin, Adam W. Feinberg

## Abstract

Microfluidic and organ-on-a-chip devices have improved the physiologic and translational relevance of in vitro systems in applications ranging from disease modeling to drug discovery and pharmacology. However, current manufacturing approaches have limitations in terms of materials used, non-native mechanical properties, patterning of extracellular matrix (ECM) and cells in 3D, and remodeling by cells into more complex tissues. We present a method to 3D bioprint ECM and cells into microfluidic collagen-based high-resolution internally perfusable scaffolds (CHIPS) that address these limitations, expand design complexity, and simplify fabrication. Additionally, CHIPS enable size-dependent diffusion of molecules out of perfusable channels into the surrounding device to support cell migration and remodeling, formation of capillary-like networks, and integration of secretory cell types to form a glucose-responsive, insulin-secreting pancreatic-like microphysiological system.

**One-Sentence Summary:** Multi-material FRESH 3D bioprinting of microfluidic CHIPS to generate fully biologic centimeter-scale and vascularized pancreatic-like tissue systems.

## Main Text

Microfluidics have rapidly advanced into cellularized and perfused organ-on-a-chip and microphysiological systems that can model increasingly complex biological processes. Important advances include lung-on-a-chip, miniature cardiac pumps, multi-organ systems, and 3D vascular networks (*1–4*). Key to this is the fluidic control, which mimics aspects of the capillary network in native tissue to provide nutrient transport and control complex interactions between different cell and tissue types. However, the polydimethylsiloxane (PDMS) silicone elastomer, thermoplastics, and photoresins used in current microfluidics inherently limit the biological relevance (*5–7*) and translation potential of these devices (*8–10*). Specifically, PDMS and thermoplastics are orders-of-magnitude stiffer than native tissue, typically absorb lipophilic biomolecules out of the media, cannot be remodeled by cells into more complex structures, and can only be used in vitro (*11, 12*). Fabricating microfluidics from other materials such as hydrogels has been recognized as a means to overcome many of these challenges, with the ability to be cellularized and better mimic the native extracellular matrix (ECM) (*5, 6, 13*). However, hydrogels remain difficult to form into complex 3D structures and the soft lithography techniques used by most studies produce fairly simple 2.5D tissue designs. Thus, a primary obstacle is developing a fabrication process for microfluidic devices which can expand both the biomaterials and cells that can be used, as well as the 3D complexity of the model systems that can be built.

Here we report the direct 3D bioprinting of collagen-based hydrogels, ECM and cells into fully-biologic microfluidic devices with high-fidelity control of structure and composition. Specific advantages include rapid fabrication and design iteration, high spatial resolution, true 3D control of device architecture, integration of multiple cell types through direct printing and perfusion post-fabrication, and integration with custom-designed vasculature and perfusion organ-on-a-chip reactor (VAPOR) bioreactors. These microfluidic devices remodel into cellularized constructs with multi-scale vascular-like networks that can develop physiologic function, demonstrated here for perfusion culture of a glucose-responsive, insulin-secreting microphysiological system. To do this we leverage freeform reversible embedding of suspended hydrogels (FRESH) 3D bioprinting (*14, 15*) to combine additive manufacturing of microfluidics with cell-mediated ECM remodeling and morphogenesis to bridge the macro (>1 mm) and micro (<1 mm) length scales. These devices are FRESH 3D bioprinted from cells, collagen type I, fibrin, and other ECM components and growth factors into complex 3D geometries termed collagen-based high-resolution internally perfusable scaffolds (CHIPS). Though high-resolution 3D printing of microfluidics using photoresins such as polyacrylates (*16*) and biocompatible hydrogels like PEG-diacrylate and gelatin methacryloyl (*4*) are established, these approaches are typically limited to a single material and at least some of the cells must be perfusion seeded after device fabrication (*17*). Similarly, extrusion 3D bioprinting of vascular-like networks using sacrificial polymers can produce perfusable tissue constructs, however, the spatial resolution and design complexity are limited (*18, 19*). Our approach has the advantages of multi-material extrusion-based 3D printing to combine multiple cell-laden and ECM hydrogel bioinks into integrated 3D structures together with the high-resolution that previously could only be achieved with light-based 3D printing. Further, as the primary structural protein in the body providing mechanical strength as well as defining vascular and tissue compartments (*20*), collagen type I is an ideal material to serve as the structural bioink for CHIPS. Together, the microfluidic CHIPS and VAPOR bioreactor form an integrated system that can recreate established microfluidic capabilities while enabling a new class of devices that combine cells with ECM proteins such as collagen and fibrin into functional, tissue-scale systems.

## Results & Discussion

### Fabrication of CHIPS via FRESH 3D Bioprinting

The CHIPS are fabricated using FRESH 3D bioprinting of collagen-based bioinks as the main structural component, replacing PDMS, photoresins or thermoplastics as the main material traditionally used in microfluidics. Collagen hydrogels used in the literature typically range in concentration from 3-10 mg/mL, which are quite soft and well below the concentration of 40-80 mg/mL found in human tissues (*21*). Using FRESH we overcome this limitation to 3D bioprint collagen bioinks with concentrations up to 35 mg/mL, resulting in constructs that can be handled and maintain shape fidelity. In FRESH, bioinks are printed in a thermo-reversible support bath consisting of a gelatin microparticle slurry designed to trigger rapid gelation during extrusion (Fig. 1A) (*14, 15*). The aqueous fluid phase of the support bath is pH-buffered to rapidly neutralize the acidified collagen bioink and drive the self-assembly of a fibrillar network, while the Bingham-plastic rheology of the support bath provides mechanical support for the embedded collagen filaments. Cell-laden fibrinogen bioinks are FRESH printed in a similar manner, where thrombin in the support bath enzymatically triggers fibrin gelation during extrusion (*15, 22*). Upon print completion, the support bath is melted by raising the temperature to 37°C, allowing for non-destructive print retrieval.

**Fig. 1.**
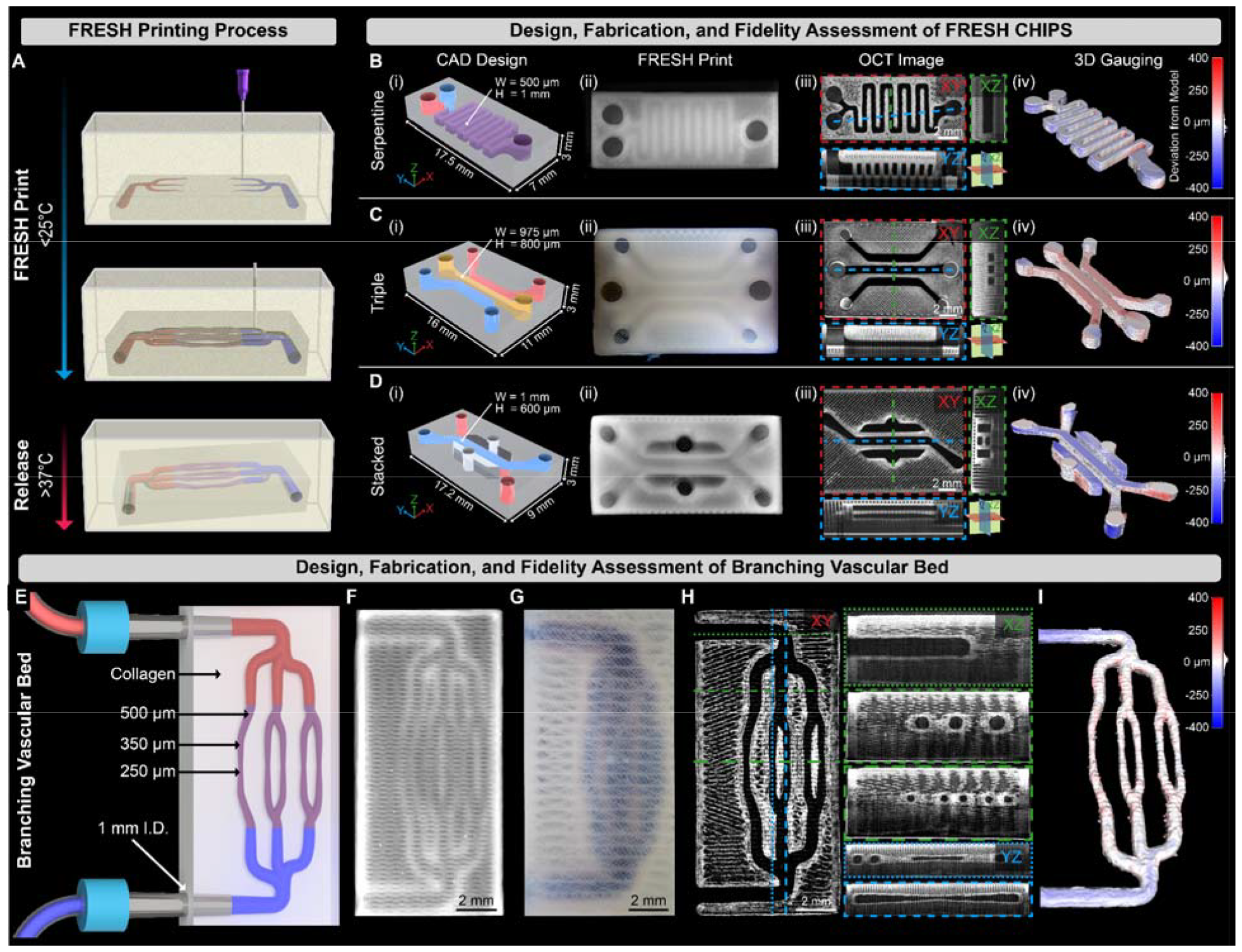
Fabrication, 3D gauging and perfusion of FRESH printed CHIPS. (**A**) Schematic of the FRESH bioprinting and release process. (**B** to **D**) CAD design (i), FRESH printing using a collagen bioink (ii), volumetric OCT imaging revealing patent lumens (dark regions, (iii), and 3D gauging of the luminal region for quantitative fidelity assessment of Serpentine, Triple, and Stacked channel internal network designs. (**E**) A multi-scale vascular bed design with open lumens from 1 mm to 250 μm. (**F**) Multi-scale vascular bed FRESH printed from collagen I. (**G**) Perfusion of blue dye through the multi-scale vascular scaffold. (**H**) Volumetric OCT imaging and cross-sectional analysis demonstrating lumen patency and circular fidelity of the internal fluidic network. (**I**) 3D gauging of the multi-scale vascular lumen volume reveals high-fidelity printing with average deviations < 11 μm.

To demonstrate the ability to fabricate collagen-based microfluidic devices, we FRESH printed a range of designs from the literature commonly made from PDMS. First, we designed a serpentine mixing network using computer-aided design (CAD) software, similar to designs reported by Whitesides and co-workers, with channel dimensions 500 μm wide and 1,000 μm tall (Fig. 1B) (*23*). The translucent nature of the FRESH printed collagen enabled visualization of the microfluidic serpentine network, while 3D imaging using optical coherence tomography (OCT) provided detailed structural data throughout the 3D volume of the device. Quantitative gauging comparing the dimensions of the FRESH printed serpentine channel to the original CAD model demonstrated the excellent print fidelity achieved, with average overprint and underprint root mean squared (RMS) errors of less than 20 μm. Next, we recreated a more complicated microfluidic design by Kamm and co-workers with 3 inlets and 3 outlets used to study vasculogenesis (*24*). This design highlights the ability to fabricate well-defined channels within a construct separated by narrow collagen walls (Fig. 1C). We also fabricated a stacked channel, lung-on-chip device based on a design by Ingber and co-workers (*1*), showing the ability to create a multi-layer design with channels in different Z-planes using a one-step FRESH printing process (Fig. 1D). Finally, we designed a vascular-like network with circular lumens branching from 1 mm inner diameter (I.D.) inlets down to 250 μm I.D. (Fig. 1E). This was FRESH printed with high fidelity (Fig. 1F) and manually perfused with a blue dye to demonstrate network patency and integrity of the channel walls (Fig. 1G). Quantitative OCT-based 3D analysis confirmed patent channels, clearly-defined individual collagen infill features outside the channels, and circular channel cross-sections (Fig. 1H). The average overprint and underprint RMS error of the channels was 11 μm (Fig. 1I), showing that FRESH printed CHIPS have excellent fidelity for channels that have both square (Fig. 1, B to D) and circular (Fig. 1, E to I) cross-sections. Together, these examples demonstrate that fully biologic microfluidic CHIPS can be designed and FRESH 3D bioprinted in a one-step process, whereas traditional microfluidics require photomask creation, sequential photolithographic mold fabrication and assembly, specialized clean room facilities, and a multi-day fabrication process (*1*).

### Microfluidic Flow within CHIPS using VAPOR Bioreactors

Microfluidic chips made from PDMS, thermoplastics and photoresins are rigid enough to support direct inlet and outlet cannulation for perfusion, but the softer FRESH 3D bioprinted collagen required a different approach to prevent leaking when pressurized. To overcome this, we developed the VAPOR bioreactor consisting of two components: (i) a main body 3D printed from UV-curable biocompatible resin with Luer lock connection fittings and internal fluidic channels, and (ii) a removable lid containing a silicone gasket and glass imaging window (fig. S1, A to D). The internal fluidic channels terminate in barbed fittings, a critical design feature, as all CHIPS for VAPOR have the negative of the barb incorporated into the inlet and outlet geometry to form an interlocking connection when assembled (Fig. 2A). A peristaltic pump, media reservoir, and bubble trap complete the perfusion circuit (fig. S1E) and the entire setup is placed in a cell culture incubator for cellular perfusion studies (fig. S1F). Since both the CHIPS and VAPOR are 3D printed, the overall dimensions and placement of inlets and outlets, as well as the internal perfusable network are easily modified and iterated upon to meet specific experimental needs.

**Fig. 2.**
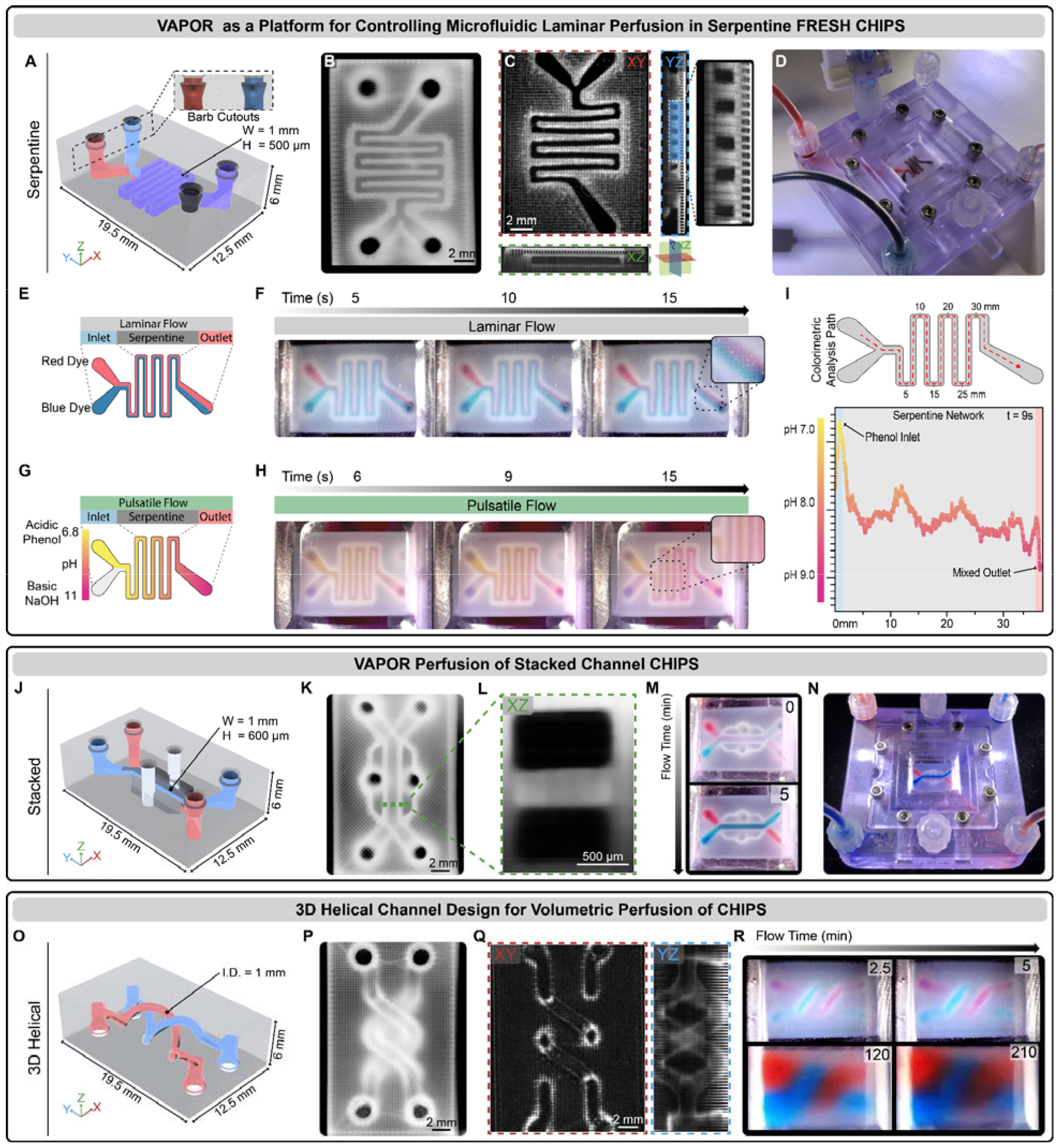
VAPOR perfusion of FRESH CHIPS. (**A**) CAD design of a dual inlet single outlet serpentine network CHIPS with barb-shaped cutouts for mating to the VAPOR inlets and outlets. (**B**) Stereomicroscope image of a FRESH printed serpentine CHIPS. (**C**) OCT volumetric imaging and cross-sectional analysis revealing patent 500 μm lumens (dark regions). (**D**) VAPOR perfusion of Serpentine CHIPS with red and black dye. (**E**) Graphic illustration of a laminar flow perfusion experiment to demonstrate balanced laminar flow of two fluids within a serpentine CHIPS. (**F**) Serpentine CHIPS perfused with a red and blue dye demonstrating laminar perfusion and minimal interfacial mixing. (**G**) Graphic illustration of a pulsatile flow perfusion experiment to demonstrate fluidic mixing within a serpentine network using an acidic phenol red indicator dye and basic NaOH solution. (**H**) Pulsatile flow in Serpentine CHIPS produces a mixing gradient along the fluidic path. (**I**) Quantitative colorimetric analysis of the pH-based mixing profile achieved within the Serpentine channel as the phenol red is neutralized. (**J**) CAD design and adaptation of Stacked channel CHIPS for interfacing with VAPOR. (**K**) Stereomicroscope image of a FRESH printed Stacked CHIPS. (**L**) An OCT XZ cross-sectional image of the stacked channels highlighting the fidelity and resolution achieved via FRESH bioprinting collagen. (**M**) Images showing initiation of Stacked CHIPS perfusion of red and blue dye in VAPOR. (**N**) Overview image of the perfused Stacked CHIPS within VAPOR. (**O**) CAD model of the 3D Helical CHIPS network. (**P**) Stereomicroscope image of a FRESH-printed 3D Helical CHIPS. (**Q**) OCT volumetric imaging and cross-sectional analysis showing patent circular lumens (XY) and a YZ projection image to view the full helical network. (**R**) VAPOR perfusion of 3D Helical CHIPS for 210 min resulting in complete volumetric diffusion of low molecular weight dyes (red, blue) throughout the CHIPS.

To assess microfluidic function, a CHIPS design containing an internal serpentine channel 500 μm wide and 1 mm tall (Fig. 2A) was FRESH printed (Fig. 2B, fig. S2, A to D), verified for lumen patency by OCT (Fig. 2C, fig. S2E), placed in the VAPOR bioreactor, and then perfused (Fig. 2D). For validation of laminar flow within the CHIPS device, we simultaneously perfused red and blue dyed solutions into the inlets (Fig. 2E, Reynolds Number, Re = 0.37). Video time-lapse imaging revealed two distinct parallel dye streams throughout the serpentine channel with minimal mixing at the dye interface due to the relatively short path length (Fig. 2F). Next, we designed a pulsatile perfusion experiment to drive non-laminar mixing between an acidic phenol red solution (starting as a visible yellow solution, pH 6.8) and a clear basic NaOH solution (pH 11) (Fig. 2G). Pulsatile flow resulted in mixing and a pH-dependent color change from yellow to magenta along the length of the serpentine path (Fig. 2H). Colorimetric analysis of the phenol red pH indicator confirmed a pH change from 6.8 to 9 along the channel, with a wave-like pH profile likely due to changes in the fluid dynamics that occur at turns within serpentine networks matching observations similarly described in the field (Fig. 2I) (*25*).

We sought to further demonstrate the versatility of CHIPS by fabricating and perfusing multi-layered and helical microfluidic designs. A multi-layered CHIPS with stacked channels based on the lung-on-chip (Fig. 1D) was designed with two 1000 × 600 μm channels separated in the Z-axis by a 400 μm collagen barrier (Fig. 2, J to L, fig. S2F). Perfusion of red and blue dye within the stacked channels resulted in two independent fluid streams separated by the collagen barrier (Fig. 2, M and N). This demonstrates that the collagen walls can prevent the mixing between different fluidic channels, though at longer time scales it is expected that there will be diffusion out of the channels. For this purpose, we designed a CHIPS containing two circular channels in a double helix configuration (Fig. 2O), a design that previously could only be created in hydrogels using DLP 3D bioprinting(*4*). This was FRESH printed with high fidelity (Fig. 2, P and Q, fig. S2F) and perfused to demonstrate time-dependent diffusion of low molecular weight (∼700 Da) red and blue dyes into the bulk collagen scaffold over 3.5 hours (Fig. 2R). The diffusion observed throughout the helical CHIPS is due to a combination of diffusion through the collagen walls of the channels and diffusion through the internal open lattice structure defined by the percentage of infill during the G-code slicing. Regardless of whether CHIPS contain planar, 2.5D, or fully 3D internal channel networks, the single-step FRESH printing approach allows a wide range of CHIPS designs to be fabricated to support perfusion.

### Molecular Weight-Dependent Diffusion Within Perfused CHIPS

To further assess molecular diffusion within CHIPS, a dual parallel channel design separated by a collagen wall (Fig. 3A) was FRESH printed, verified by OCT (Fig. 3B), and then perfused within the VAPOR bioreactor (Fig. 3C). Laminar flow (Re = 0.55) and channel integrity were confirmed by perfusion and tracking of fluorescent microbeads within the channels (Fig. 3D). We observed the expected laminar flow, with a mean speed of 0.91 ± 0.30 mm/s (theoretical value calculated as 1.2 mm/s for the center of the channel). Next, we mimicked diffusion of biomolecules of different molecular weights using FITC-conjugated fluorescent dextrans from 3 to 70 kDa in one channel as a source with the opposing channel perfused with 1X phosphate buffered saline (PBS) as a sink (Fig. 3E). Time-lapse fluorescence imaging of defined regions of interest (ROIs) expanding outward from the dextran-perfused channel (Fig. 3F, fig. S3, A and B) established that diffusion throughout the CHIPS was molecular weight-dependent (Fig. 3G). The 3 kDa dextran reached steady state diffusion 1 mm (ROI 1) from the source channel within 3 hours, while the 70 kDa dextran reached a steady state rate at ∼20 hours. The diffusivity of larger 40 and 70 kDa molecules is notable compared to cast solid hydrogel scaffolds (*26*), where diffusion has been reported to be much lower. To increase molecular diffusion and mimic the interstitial pressure found in tissues, we introduced a 5 or 10 mmHg pressure differential between the dextran source and PBS sink channels (Fig. 3H) (*27*). Spectrophotometric analysis of the PBS sink perfusate after 24 hours showed significantly more 40 kDa dextran present in the perfused PBS sink following a pressure increase of either 5 or 10 mmHg (Fig. 3I). Comparing the ratio of fluorescence intensity between high pressure (HP, + 10 mmHg) and normal pressure (NP) over time showed that increasing pressure drove the dextran into the PBS channel as well as throughout the CHIPS and into the furthest distal regions (Fig. 3, J and K, fig. S3C). Together, these results establish that molecules with a wide range of molecular weights can diffuse out of the CHIPS channels and into the bulk of the construct, something that is not possible with the vast majority of microfluidic devices made from PDMS or plastics.

**Fig. 3.**
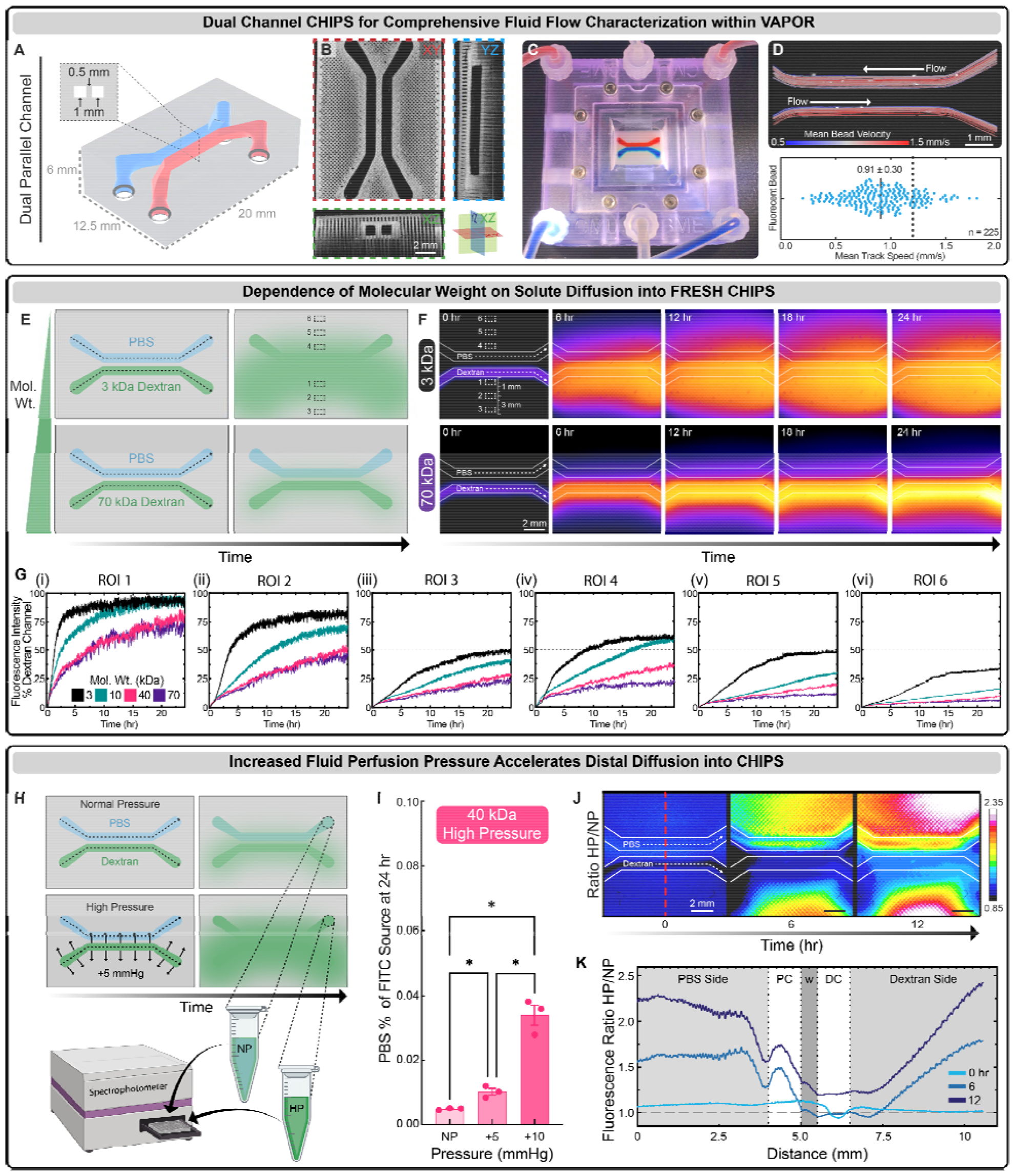
Characterization of diffusivity within FRESH CHIPS. (**A**) A CAD model of dual parallel channel CHIPS with the channels separated by a collagen wall. (**B**) OCT cross-sectional views of dual parallel channel CHIPS. (**C**) VAPOR perfusion of dual parallel channel CHIPS with red and blue dye. (**D**) Perfusion and tracking of fluorescent microbeads to measure velocity profiles and mean bead speed compared to the theoretical estimation (dotted line). (**E**) A schematic demonstrating the perfusion of dual parallel channel CHIPS where one channel i always perfused with PBS while the other channel is perfused with different sizes of FITC-conjugated dextran with 6 ROIs being selected for diffusivity analysis. (**F**) Time lapse fluorescence images displayed with fire intensity look up table of dual parallel channel CHIPS undergoing VAPOR perfusion with FITC-conjugated dextrans ranging from 3 to 70 kDa. (**G**) ROI-based Analysis of dextran fluorescence intensity over time beginning proximal to the dextran source channel (i) and ending distal to the PBS source channel (vi). (**H**) A schematic demonstrating increased pressure (+5 mmHg) within the dextran source channel driving diffusion into the perfusate from the PBS circulation that was sampled for spectrophotometric analysis. (**I**) Analysis of 40 kDa FITC-conjugated dextran fluorescence intensity within the PBS circulation after 24 hours of either normal (NP), or increased pressure (+5 mmHg, +10 mmHg) within the dextran source channel. (**J** and **K**) Timelapse images and line profile analysis of fluorescence intensity at 0, 6 and 12 hours calculated as a ratio of the high pressure (HP) image divided by the normal pressure (NP) image to reveal pressure-dependent regions of increased diffusion.

### Multi-material FRESH Printing to Create Spatially Patterned and Cellularized CHIPS

The spatial patterning of cells and ECM within internal regions of microfluidic devices is one of the major challenges in engineering more sophisticated microphysiological systems. To address this, we implemented multi-material FRESH printing using a custom 3D bioprinter with three Replistruder syringe pump extruders for bioink deposition (fig. S4) (*22, 28*). For printing validation, we designed a parallel plate CHIPS device containing an open rectangular channel to facilitate confocal imaging of the patterned biomaterials (fig. S5). Collagen bioinks doped with fluorescently-labeled fibronectin (Fn) were printed along the channel lumen to form regions ∼200 μm thick to demonstrate control over spatial patterning of ECM composition. Using multi-material FRESH printing, we created a uniform layer across the entire channel (fig. S5A), a pattern of lines 1 mm wide running the length of the channel (fig. S5B), or a defined branching pattern with segments down to 500 μm in width within the channel (fig. S5C). Finally, we patterned softer and stiffer regions along the length of the channel by printing collagen bioinks with concentrations of 6, 12, and 23 mg/mL, each labeled with different fluorescent dyes (fig. S5D).

Next, we sought to implement multi-material FRESH printing to pattern cells and ECM in 3D, with the goal to support endothelial cell attachment within our dual channel CHIPS. The ability of vascular cells to attach, proliferate, and form a network within our devices is one of the key advantages of CHIPS as a way to provide nutrient delivery and waste removal to a larger tissue volume. For our design, we choose to line the channels with a layer of 12 mg/mL collagen with fluorescently labeled Fn to improve endothelial cell attachment combined with a central region between the two channels consisting of 23 mg/mL collagen with fluorescently labeled Fn and vascular endothelial growth factor (VEGF), intended to promote cell infiltration (fig. 6A). The CHIPS was FRESH printed, optically cleared, and imaged via confocal fluorescence microscopy for visualization of the internally patterned structures (fig. S6B). The 12 mg/mL collagen and Fn channel linings were confirmed to be patterned as intended, while the central region containing VEGF was spatially restricted between the channels as designed (fig. S6C). To test cell adhesion and growth within the multi-material CHIPS, human umbilical vein endothelial cells (HUVECs) were perfusion seeded into the dual channel CHIPS at a high cell density (fig. 6D). Following 5 days of static culture, the HUVECs spread along the channel lumen, formed a visible monolayer, and stained positive for CD-31 (fig. S6, E to I). While there was some evidence of cell migration into the bulk scaffold, the HUVECs were primarily restricted to the channels into which they were seeded, potentially due to the absence of a stromal cell population present within our CHIPS (*29*).

To expand the complexity of the CHIPS we can engineer, we sought to fully leverage our multi-material printing by printing cell-laden bioinks and avoid the laborious process of perfusion seeding of cells that is required for nearly all organ-on-chip and microphysiological systems. To do this, we directly printed a vascular bioink within the channel walls to encourage HUVEC vasculogenesis followed by angiogenic sprouting. The process for printing cells with FRESH is similar to that of collagen, except rather than using a pH-change to initiate bioink gelation, thrombin is used to enzymatically trigger the cross-linking of fibrinogen into fibrin. Importantly, both collagen and cell-laden fibrinogen bioinks are FRESH printed into the same support bath as their gelation mechanisms are orthogonal to one another, enabling the 3D bioprinting of fully integrated, multi-material cellularized CHIPS. Our cell-laden fibrinogen vascular bioink contained HUVECs and human bone marrow-derived mesenchymal stem cells (MSCs) as the pericyte-like cell to support microvessel formation and stability (*30*). We evaluated multiple designs for these cellularized CHIPS, starting with a vascular bioink that was printed as both the channel lining and dividing region between dual parallel fluidic channels, while the remaining structure was collagen (Fig. 4A). Confocal imaging of the CHIPS immediately after printing confirmed our ability to volumetrically pattern both the cellular and collagen bioinks in 3D while maintaining patent fluidic channels (Fig. 4B). However, with this CHIPS design after static culture for 8 days, we observed buckling due to cell-driven compaction forces (fig. S7A). To prevent these large-scale deformations and maintain proper fit for use in the VAPOR bioreactor, we added collagen walls between the channel linings and central region as mechanical reinforcement (Fig. 4C). Multiphoton and second harmonic generation imaging of the collagen-reinforced CHIPS revealed high fidelity cellular channel linings and the presence of the extra collagen walls (Fig. 4D). After 8 days of static culture, the mechanical reinforcement provided by the collagen walls was sufficient to prevent the CHIPS from buckling compared to the fibrin-wall CHIPS (fig. S7B). These results highlight the importance of both CAD design and material properties for long-term success and performance of cellularized CHIPS and our ability to rapidly make such changes.

**Fig. 4.**
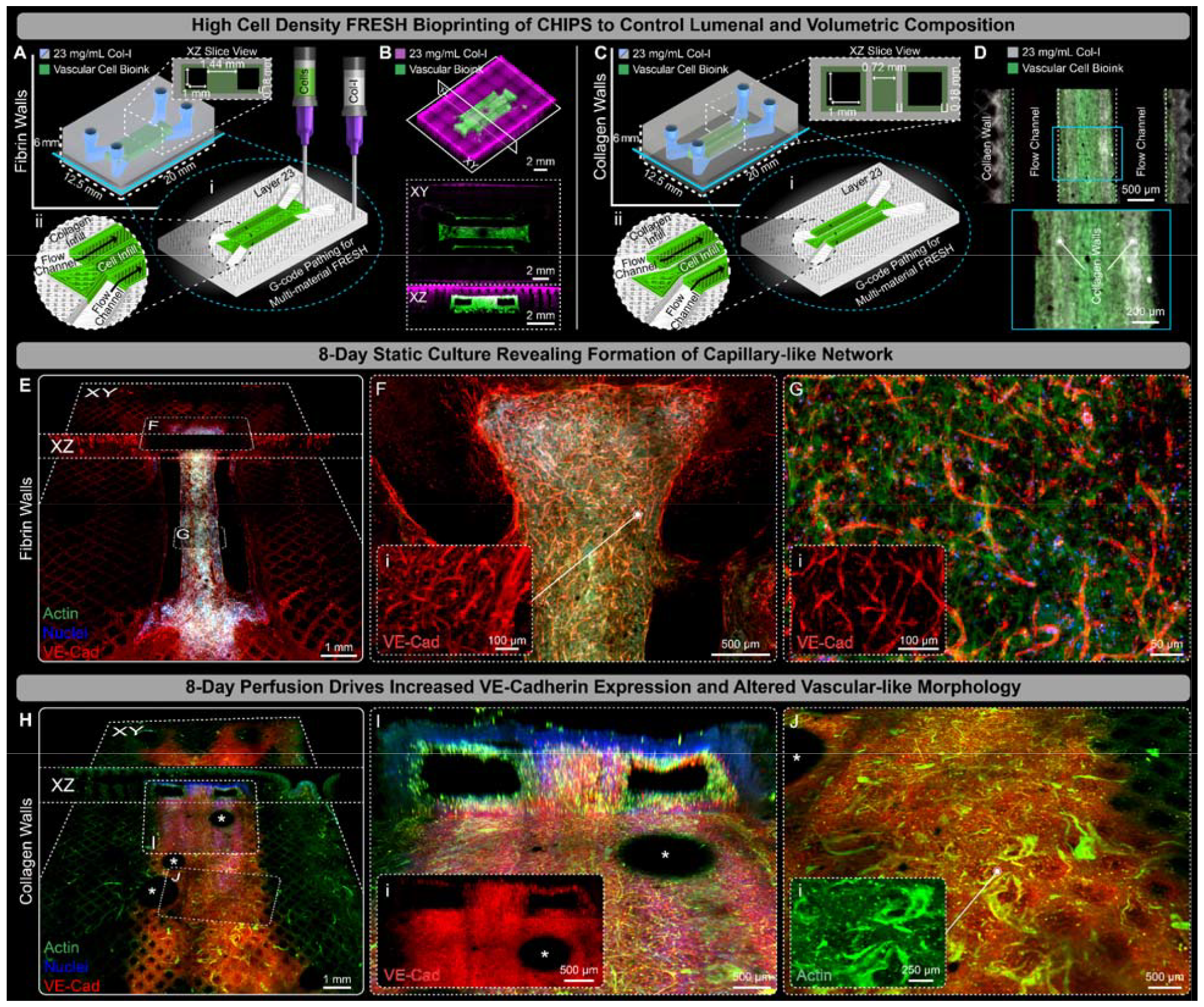
Multi-material printing of vascular cell bioink drives capillary-like network formation and enhanced VE-cadherin expressing within VAPOR perfused CHIPS. (**A**) Schematic and 3D printer machine pathing (G-code) of a dual parallel channel multi-material CHIPS with an internal fibrin-based vascular cell bioink (HUVEC and MSC) region containing 1 mm^2^ fluidic channels. (**B**) 3D confocal fluorescence imaging of optically cleared high-fidelity collagen (magenta) and vascular cellular bioink (green) printing with XY and XZ plane views revealing internal structure, open channels, and high-fidelity volumetric registration between the collagen and vascular bioink. (**C**) Schematic design and pathing G-code of dual channel CHIPS with collagen walls between the cellular channels. (**D**) An XY mid-plane view of whole mount confocal fluorescence images of optically cleared 1-day statically cultured vascular CHIPS with collagen walls. Combined multiphoton fluorescence and second harmonic imaging of the vascular bioink and collagen shows the spatial alignment between the cellular biomaterial (green) and collagen walls (gray). (**E**) Both XY and XZ midplane slice views from 3D confocal imaging of optically cleared dual parallel channel cellular CHIPS with fibrin walls statically cultured for 8 days and stained for actin (green), nuclei (blue), and the endothelial marker VE-Cadherin (VE-Cad, red). (**F** and **G**) Confocal images of regions within (E) revealing microvascular cell spreading and migration (F), and vascular network formation (G) with inset images displaying magnified views of the VE-Cad only channel. (**H**) Both XY and XZ slice views of the channel bottom surface from 3D confocal imaging of optically cleared dual parallel channel cellular CHIPS with collagen walls following 8 days of perfusion culture within VAPOR. (**I** and **J**) Confocal images revealing strong lumenal expression of VE-Cadherin (I), and evidence of cellular remodeling and vascular-like network maturation (J) with inset images displaying magnified views of the VE-Cad and Actin only channels respectively. *Indicates air bubble artifacts introduced into the channels during CHIPS optical clearing and imaging.

Having demonstrated that vascular cells can be directly FRESH printed within CHIPS, we investigated the importance of culture time and perfusion on cell behavior. After 8 days of static culture in CHIPS with fibrin walls, large areas of vascular endothelial cadherin (VE-Cad)-expressing microvascular-like structures were observed within and adjacent to the cellular printed regions (Fig. 4E). Despite the absence of perfusion, the cells appeared to form networks spanning distances >2 mm, migrate from the ends of the printed cellular regions into the acellular collagen (Fig. 4F), and established dense, microvascular-like networks throughout the central dividing region (Fig. 4G). We observed similar results after 8 days of static culture in CHIPS with collagen walls (fig. S8A), with VE-Cad expression along the open channel lumen (fig. S8B), and migration of cells (>1 mm) beyond the edge of the printed cellular regions from where the cells originated (fig. S8C). Importantly, this was the first evidence we found that cells can preferentially migrate along collagen filaments. While static culture conditions can result in capillary-like network formation, the addition of perfusion has been shown to increase endothelial cell-specific protein expression, and promote angiogenesis (*31*). After 8 days of VAPOR perfusion culture in CHIPS with fibrin walls (fig. S8D) we observed high levels of VE-Cad expression encircling the printed channels (fig. S8E) and cell elongation within the central dividing region (fig. S8F). The results in the CHIPS with collagen walls after 8-day perfusion were comparable, with extensive cell migration throughout the device and formation of capillary-like structures far from where the HUVECs and MSCs were printed (Fig. 4H). Vascular CHIPS with collagen walls were more stable in the VAPOR bioreactors during perfusion, maintaining larger channel openings that remained lined with cells (Fig. 4I). Furthermore, evidence of perfusion-stimulated changes of the HUVECs and MSCs into a vascular-like morphology reminiscent of larger diameter microvessels was observed within the VE-Cad-rich zones all around the fluidic channels (Fig. 4J). These results demonstrate that VAPOR perfusion of vascular CHIPS promotes cell assembly into capillary-like networks spanning 8-100 μm in diameter by supporting initial vasculogenesis followed by angiogenesis throughout the scaffold.

### Pancreatic-like CHIPS Demonstrate Glucose Stimulated Insulin Response

Secretory function is a key role of many organs in the body, and we focused on integrating additional cell types into our perfused CHIPS to demonstrate the complexity and physiology that can be achieved. Specifically, we focused on glucose stimulated insulin secretion (GSIS) by islets of the pancreas, the failure of which leads to type I diabetes. The engineering of pancreatic tissue has made a number of advances, including hydrogel-based encapsulation to evade the immune system (*32, 33*) and microphysiological systems containing isolated islets (*34–36*), showing that implantation of insulin-producing engineered tissues has therapeutic potential (*34, 37*). However, there has yet to be a perfusable tissue construct created entirely out of ECM and cells with any of these approaches. To model the GSIS of pancreatic tissue, mouse MIN6 cells were chosen as they exhibit beta-like cell secretion of insulin in response to glucose. MIN6 cells were incorporated into the vascular bioink to form a high-concentration (60 million cells/mL total) pancreatic-like bioink for multi-material FRESH printing into the parallel channel CHIPS design. The first pancreatic-like CHIPS design we evaluated resembled the dual channel fibrin-wall CHIPS (Fig. 4) with the cell-laden bioink lining the fluidic channels and within the central dividing region (fig. S9, A and B). The pancreatic-like CHIPS were FRESH printed and statically cultured for 8 days where construct size remained relatively unchanged (fig. S9C). After fixing and imaging, we observed dense cellularization in the regions where cells were printed as well as migration of cells into the surrounding collagen scaffold (fig. S9D). In addition, the CHIPS exhibited patent fluidic channels and stained positive for insulin expression within both the central dividing region and adjacent channel walls (fig. S9, E to G). Within the cellularized regions, we also observed a dense, capillary-like network staining positive for actin and CD-31 (fig. S9H). There was also further evidence that we can guide cells to align along the direction of the printed collagen filaments, as cells appeared to follow the 45° surface filament pattern of the printed collagen (fig. S9, I and J). Cells were also seen migrating up to 200 μm from the cellular regions into the collagen scaffold along the printed collagen infill filaments (fig. S8, K and L), indicating that the topology of printed filaments can serve as a potential pathway to define network density and branching.

While the static pancreatic-like CHIPS did express insulin, a large volume of our construct remained acellular. In principle, since our entire scaffold is fabricated from ECM, we can increase the total cellular volume within the CHIPS by printing additional cellularized regions. A modified pancreatic-like CHIPS was designed with three distinct cellularized regions between and adjacent to each side of the channels to increase the printed cell volume by 50%, and included printed collagen channel linings to strengthen the device and prevent cell-mediated deformation (Fig. 5A). The CHIPS were successfully FRESH printed and after static culture for 12 days displayed extensive cell migration into adjacent acellular regions (Fig. 5B). Notably, the channel lining that was initially acellular collagen now had a lumen with dense capillary-like networks that bridged the original printed regions (Fig. 5C, fig. S10A). In addition to the cell migration into and around the fluidic channels, extensive cell migration outward into the surrounding collagen scaffold was observed a full millimeter from the initial printed regions (fig. S10, B and C). The channel lumens remained patent and the originally acellular walls were now densely populated with cells (fig. S10D). Analysis of the segmented branching network growing into the channels revealed a mean diameter of 9.3 ± 2.7 μm (Fig. 5C). Cells were also observed migrating hundreds of microns into the scaffold along the printed collagen filaments (Fig. 5D).

**Fig. 5.**
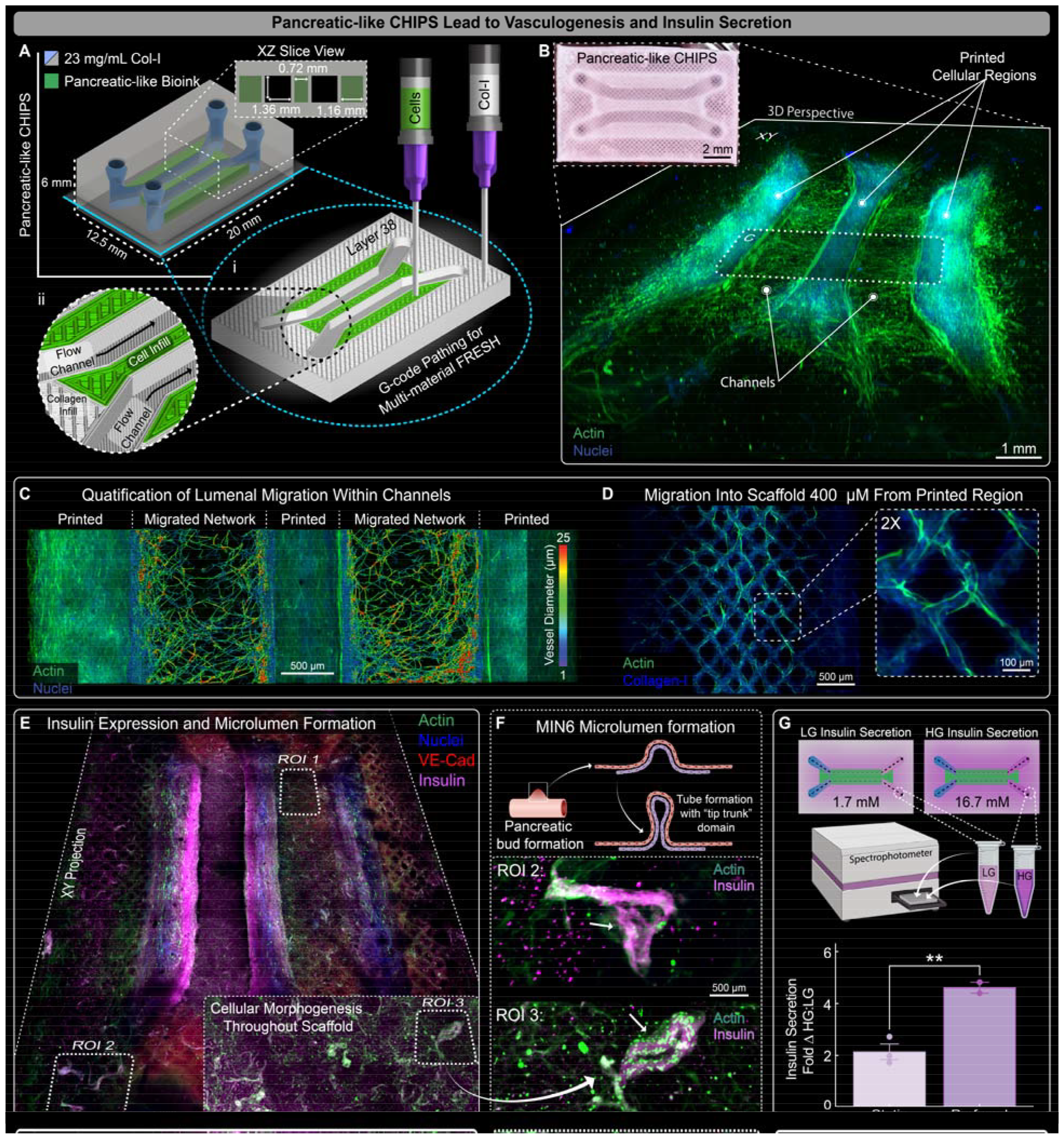
FRESH printed vascular pancreatic CHIPS demonstrate vasculogenesis and insulin secretion. (**A**) Schematic design and 3D printer machine pathing G-code of a dual parallel channel multi-material CHIPS with pancreatic vascular cell bioink (MIN6, HUVEC, and MSC) regions surrounding both sides of the channels. (**B**) Pancreatic CHIPS FRESH printed and visualized via brightfield stereomicroscope (inset) and 3D confocal fluorescence imaging of the optically cleared pancreatic scaffold after 12 days of static culture. (**C**) XY midplane view from confocal fluorescence image of the 12-day statically cultured pancreatic CHIPS following 3D vascular network segmentation for quantification of network diameter and density within the migratory zones. (**D**) Example confocal fluorescence images revealing additional cell migration into the acellular regions of the CHIPS beneath the cellular regions guided by the printed collagen filaments. (**E**) XY midplane projection view from 3D confocal fluorescence imaging of the optically cleared 12-day VAPOR perfused pancreatic CHIPS. (**F**) Graphic illustration and example regions of interest (ROIs) depicting evidence of early MIN6 pancreatic bud and microlumen formation with actin (green) and insulin (magenta) fluorescence images from ROI 2 and 3 in (E). (**G**) Graphic illustration and quantification for insulin secretion ELISA assay from 1.5-hour glucose-stimulated (ratio of high glucose-HG to low glucose-LG) insulin secretion experiment between 12-day static and perfusion cultured pancreatic CHIPS (mean + STDEV; **P < 0.01 for n= 2 static tissues; n = 2 perfused tissues).

To improve nutrient delivery, increase cell proliferation and growth, and enhance MIN6 insulin secretion, we perfused the pancreatic-like CHIPS in the VAPOR bioreactor for 12 days. Perfusion resulted in increased cellularization throughout the device, with vascular-like structures ranging in diameter from 25-100 μm originating from the printed cellular regions and following the printed collagen filaments (Fig. 5E, fig. S11, A to D). We also found extensive insulin expression within all printed cellular regions along with insulin-positive structures far from the printed cellular regions, suggesting co-migration of the MIN6 cells as well as the HUVECs and MSCs that formed the capillary-like structures (Fig. 5E). Indeed, one of the most exciting observations was the morphogenesis of looping structures that contained insulin-positive MIN6 cells and actin-positive HUVECs and/or MSCs, reminiscent of early branching morphogenesis observed during pancreatic islet development (Fig. 5F) (*38–40*). To confirm that the MIN6 positive insulin staining was indicative of functionally secreted insulin in response to glucose stimulation, we performed a GSIS ELISA assay comparing the static and perfusion-cultured pancreatic-like CHIPS. Note that due to the thickness of the pancreatic-like CHIPS (∼6 mm) and resulting insulin diffusion time, we were required to restrict our experiment to one low and one high glucose stimulation per device without a second low glucose stimulation. Results showed that pancreatic-like CHIPS cultured statically produced a 2-fold increase in insulin secretion following glucose stimulation whereas those cultured under perfusion exhibited a statistically significant 4.6-fold increase (Fig. 5G). This places the performance of pancreatic-like CHIPS between that reported for MIN6 pseudo-islet spheroids and primary human islets, which frequently produce stimulation indices of approximately 1.5 to 3-fold and 10-fold, respectively (*34–36, 41–45*). Further, we quantified the amount of insulin secreted over 24 hours of perfusion after a fresh media exchange. We detected >8 ng of additional insulin present within the media perfused through the pancreatic-like CHIPS compared to the control media sample. This result demonstrates that placing beta cell-like MIN6 in a fully biologic 3D ECM environment with appropriate vascular cell types and perfusion can drive the formation of a tissue construct with sustained secretory function. Beyond these proof-of-concept studies for the pancreatic-like CHIPS, further functional improvement is likely achievable through multiple approaches including increasing the beta-like cell volume within the construct, extending the culture time to promote more mature cell phenotypes, and incorporating fully differentiated human iPSC-derived beta-like cell clusters or human primary islets.

## Conclusion

In summary, we have FRESH 3D bioprinted microfluidic CHIPS made entirely from cells and ECM proteins instead of traditional materials such as PDMS, opening up a range of new capabilities for organ-on-a-chip and microphysiological systems. While PDMS, photoresins and thermoplastics are staples for microfluidic chip production, there are fundamental material limitations that restrict their use in developing more advanced applications that require a fully biologic environment. The inability of PDMS and these other polymers to undergo cell-driven remodeling restricts the complexity to casting cellularized hydrogels within channels (*27*) or seeding them on luminal surfaces (*46*). In clear contrast, we show that by FRESH printing CHIPS from ECM proteins, we allow for cell-driven remodeling and migration broadly throughout the device into regions that would typically be inaccessible within traditional microfluidics. Further, perfusion with the VAPOR bioreactor supports tissue viability, long-distance cell migration within the CHIPS, self-assembly of capillary-like networks, and the emergence of tissue-scale function as demonstrated by insulin secretion in the pancreatic-like CHIPS. We also believe this approach will create a new type of engineered tissue construct that blurs the line between in vitro and in vivo systems. While microfluidics has provided a powerful platform for observing tissue morphogenesis, disease mechanisms, and pharmaceutical response (*47*), the inert nature of PDMS and other polymers precludes their suitability for long-term *in vivo* implantation. This is where *in vitro* maturation of fully biologic CHIPS via VAPOR bioreactor perfusion provides a path forward for generating larger, pre-vascularized tissues with improved viability, functionality, and therapeutic potential upon implantation. For example, the human pancreas contains ∼1 billion beta cells(*48*), which using our current pancreatic-like bioink of 30 million MIN6/mL would require an ∼33 cm^3^ construct to match cell number. However, we have previously demonstrated FRESH printed bioinks containing >200 million cells/mL (*15, 22*), which would reduce the pancreatic-like bioink volume to ∼5 mL, only ∼3 times larger than our current CHIPS. This suggests that in the future it may be possible to engineer CHIPS that can recreate at least some aspects of full organ-scale function, whether as in vitro devices for disease modeling and pharmacology, or as in vivo devices designed for implantation as a cell replacement therapy.

## Author contributions

Conceptualization: DJS, ARH

Methodology: DJS, ARH

Investigation: DJS, ARH, JT, EB, SM, BDC

Visualization: DJS, ARH, AWF

Funding acquisition: DJS, AWF

Project administration: DJS, AWF

Supervision: DJS, ARH, AWF

Writing – original draft: DJS, ARH, EB, SM, BDC AWF

Writing – review & editing: DJS, ARH, EB, SM, BDC AWF

## Data and materials availability

All data are available in the main text and the supplementary materials. The STL files for 3D bioprinter hardware modifications, collagen constructs, and perfusion systems are available under an open-source CC-BY-SA license at https://3dprint.nih.gov/users/awfeinberg.

## Supplementary Materials

Materials and Methods

Figs. S1 to S11

References (*49-55*)

