## Supplementary Materials for "3D Bioprinting of Collagen-based Microfluidics for Engineering Fully-biologic Tissue Systems"

**The PDF file includes:**
Materials and Methods
Figs. S1 to S11
Captions for Movies S1 to S10
References

**Other supplementary material for this manuscript includes:**
Movies S1 to S10

### Materials and Methods

3D Bioprinter Setup

All FRESH printing was executed on custom-built 3D bioprinters utilizing our open-source Replistruder 5 syringe pumps (*22*, *28*, *49*) (parts files available upon publication for download under a CC-BY-SA license found at https://3dprint.nih.gov/users/awfeinberg). Single material printing utilized a custom gantry 3D printer configuration driven by four Parker Hannifin 404 × R 100 mm travel precision stages (8 μm travel accuracy) as previously described (*22*). For multi-material bioprinting a high-performance motion control system was designed utilizing the AGS1000 platform (Aerotech Inc.). Three Replistruder 5 syringe pumps and a dovetail mounted OCT scanhead (Thorlabs) were each mounted to TU-30 linear ball screw stages (IKO) for independent actuation of each extruder. The custom bioprinter platform resulted in a build volume of 100 mm (X), 300 mm (Y), 100 mm (Z), and 80 mm for each TU-30 substage. A 500 µL, 1 mL, or 2.5 mL syringe (Hamilton) containing bioink with a stainless-steel needle of either 80 (Jensen Global, JG34-0.25HPX) or 150 (Jensen Global, JG30-0.5HPX) µm inner diameter (ID) was used.

Collagen Bioink Preparation

Unless stated otherwise, a 23 mg/mL acidified collagen bioink was utilized for all prints and prepared as previously described (*15*). Briefly, sterile 35 mg/mL neutral collagen bioink (LifeInk 200, Advanced Biomatrix, 5278) was diluted in a 2:1 volume ratio with 0.24M acetic acid (VWR, 97064-482) or sterile 35 mg/mL acidified collagen bioink (LifeInk 240, Advanced Biomatrix, 5267) was diluted in a 2:1 volume ratio with sterile DI $H_2O$ and mixed back and forth 40 times between two mated syringes. For the preparation of fluorescent bioinks, acidified collagen bioinks were mixed with 10-20 µL of 500 µg/mL human fibronectin (Corning, 356009) fluorescently conjugated to either Alexa-Fluor 405, 488, 555, and 633 NHS Esters (*50*, *51*). In multi-material printing experiments, the final concentration of the fibronectin within the collagen bioink was 50 ug/mL. In all cases, syringes containing acellular bioink were centrifuged at 3000 g for 5 min at room temperature to remove any air bubbles generated during the preparation process. The bioink was then transferred to a Hamilton glass syringe for printing.

Cell Culture

Unless otherwise stated, cells were cultured at 37°C under 5% $CO_2$ with media supplemented with 1% (v/v) penicillin-streptomycin being exchanged every 2 days. Pooled human umbilical vein endothelial cells (HUVEC) (Lonza, CC-2519) were cultured in endothelial cell media (Lonza, EGM-2, CC-3162). Bone marrow-derived MSCs (ATCC, PCS-500-012) were cultured on flasks coated with quick coating solution with growth media (ATCC, PCS-500-041). MIN6 (Mouse insulinoma cell line) (Addexbio C0018008) were cultured in high-glucose DMEM (Gibco, 11-965-092) with 15% FBS (v/v), 2 mM sodium pyruvate, 20 mM HEPES, and 0.05 mM β-mercaptoethanol, using passages 9-15. MIN6 cells were seeded at a density of 2 X $10^4$ cells/$cm^2$ and passaged when they reached 80-90% confluency.

Cellular Bioink Preparation

Vascular and pancreatic bioinks were prepared for cellular bioprinting experiments following similar protocols. For the vascular bioink, HUVECS (passage 4-6) and MSCs (passage 2-4) were cultured following the previously described procedure and lifted using a trypsin-EDTA solution. Trypsin was neutralized using trypsin neutralizing solution with 7.5 µM bivalirudin

(Cayman Chemical, 23035) as a thrombin inhibitor at a 1:2 ratio. The cells were pelleted at 200 g for 5 minutes and then resuspended in 1 mL Hank's Balanced Salt Solution (HBSS, Gibco, 14175-095). 1 X $10^6$ MSCs and 9 X $10^6$ HUVECs were transferred into a 1 mL BD syringe. This syringe was then centrifuged at 190 g for 5 minutes and the supernatant was aspirated until approximately 100 µL remained. 165 µL of 120 mg/mL fibrinogen (Millipore Sigma, 341573) and 65 µL 5% xanthan gum (dissolved in HBSS) were loaded into a separate 500 µL Hamilton gastight syringe. The 1 mL and 500 µL syringes were connected with a female luer lock adapter and the fibrinogen, xanthan gum, and cells were mixed 50 times. The vascular bioink was centrifuged at 300 g for 3 minutes in the 1 mL syringe to remove bubbles and transferred to the 500 µL syringe. The final vascular bioink consisted of 30 X $10^6$ cells/mL, 60 mg/mL fibrinogen, and 1.0% (w/v) xanthan gum. The pancreatic bioink was prepared similarly to the vascular bioink with the following adaptations. MIN6 cells (passage 9-15) were lifted using trypsin solution for 5 min, and 10 X $10^6$ MIN6 cells were added to the 1 X $10^6$ MSCs and 9 X $10^6$ HUVECs in a 1 mL syringe. The final pancreatic bioink consisted of 60 X $10^6$ cells/mL (30 X $10^6$ MIN6/mL, 27 X $10^6$ HUVEC/mL, 3 X $10^6$ MSC/mL), 60 mg/mL fibrinogen, and 1.0% (w/v) xanthan gum.

<u>FRESH Support Bath Generation</u>

Cellularized CHIPS were printed using a sterile support bath (LifeSupport, FluidForm) prepared according to the manufacturer's instructions. When printing collagen-based bioinks the support bath was rehydrated with a 2:1 mixture of cold 100 mM 4-(2-hydroxyethyl)-1-piperazineethanesulfonic acid (HEPES), pH 7.4 (Corning, 60-034-RO) and serum free DMEM (Gibco, 11-965-092). When printing fibrinogen-based bioinks, the support bath was rehydrated with cold 100 mM HEPES, pH 7.4, and 1 U/mL thrombin (Millipore Sigma, T4648). Acellular CHIPS were printed using a FRESH support bath generated using a complex coacervation method as previously described (*15*). Briefly, FRESH v2.0 support bath (*15*) was made by dissolving 3.0% (w/v) gelatin type B (Fisher Scientific, G7-500), 0.3% (w/v) gum arabic (Sigma Aldrich, G9752), and 0.125% (w/v) Pluronic® F-127 (Sigma Aldrich, P2443) in a 50% (v/v) ethanol solution at 45°C. Soon after dissolving, the pH of the solution was adjusted to 5.65 using 1M hydrochloric acid. The solution was sealed and stirred overnight at room temperature. The following morning the slurry was centrifuged at 300 g for 2 min, the supernatant discarded, replaced with DI $H_2O$ and shaken to wash the particles. The slurry was recompacted by centrifuging at 750 g for 3 min, the supernatant discarded, and replaced with DI $H_2O$. This washing step was repeated a total of 3 times. After the final round of washing the slurry was resuspended to a final concentration of 100 mM HEPES, pH 7.4 and stored at 4°C. Prior to printing, the slurry was placed in a vacuum chamber at room temperature for 30 min followed by centrifugation at 2000 g for 5 min. The supernatant was discarded, and the slurry was transferred into the print container of choice.

<u>CHIPS Design</u>

CHIPS were created in computer-aided design (CAD) software (Autodesk Inventor; Autodesk Fusion 360). All models were exported as STL files prior to printing. Overall design features and dimensions varied between experimental conditions and requirements. Perfusable networks were designed as void space within the printed CHIPS. All STL files used in the manuscript can be found at https://3dprint.nih.gov/users/awfeinberg. 3D renders for manuscript preparation were generated within the rendering environment and exported as PNG files. Assembly videos of components were made within the animation environment and exported as .avi or .mp4 files.

FRESH 3D Bioprinting of Collagen Type I

Collagen type I was FRESH printed as previously described (*15*). All STL files were sliced using slicing software (Ultimaker, Cura; PrusaResearch, PrusaSlic3r; Slic3r) to produce G-code files. For 80 and 150 µm ID needles, a layer height of 32 and 60 µm was used, respectively. Chips were printed at 23-70 mm/s with 2 perimeters, 4 top and bottom layers, and 35% infill. All constructs were printed at room temperature (22$^{o}$C). Upon completion, constructs were incubated at 37$^{o}$C for at least 30 min to melt the support bath. The molten support bath was exchanged with warm print storage solution (PSS) consisting of consisting of 1X PBS, 50 mM HEPES, pH 7.4, and 1% (v/v) penicillin-streptomycin (Life Technologies, 15140-122). Acellular CHIPS were optionally sterilized by 10 min UV ozone treatment followed by overnight incubation in PSS to remove residual gelatin.

Brightfield and Stereoscopic Imaging

For all brightfield and stereoscopic images we utilized a Leica M165FC microscope with a 1X Plan Apo lens, fully adjustable base with darkfield capability, and a Prime 95B CMOS camera. Additionally, images of printed CHIPS and various equipment utilized were taken with either a Soney A5 camera equipped with a Laowa 24mm f/14 probe lens, or an iPhone 14 pro. Image contrast adjustment and resizing was performed in FIJI ImageJ or Adobe Photoshop.

OCT Imaging and 3D Gauging of CHIPS

The full 3D structure of CHIPS were imaged using optical coherence tomography (OCT) to non-invasively assess lumen patency for quality control and in-process monitoring as previously described (*22*). Briefly, OCT images were acquired using a Vega 1300 nm OCT system (Thorlabs, VEG210C1) mounted onto the bioprinter using an objective (OCT-LK4 objective) with an imaging depth of 11 mm and 13 µm lateral resolution. XYZ voxel sizes were acquired at 16.22 x 16.22 x 8.11 µm, respectively. Once scanned, XY, XZ, and YZ planes of the 3D images were analyzed to qualitatively assess the print for patency, significant defects, and potential blockages within the printed networks that would compromise flow through the CHIPS. Computational 3D gauging was performed using a combination of FIJI ImageJ, Imaris (Bitplane v9.5), 3D Slicer, and Cloud Compare software. Raw OCT images were denoised and scaled to account for refractive index of the imaging medium. Processed images were exported as TIFF stacks and imported to either Imaris or 3D slicer for segmentation of the internal perfusable networks (*52*). The segmented internal networks were exported as .STL files in 3D Slicer and imported into 3D builder to center the model at the origin. Both the segmented .STL and original .STL used for printing were imported into Cloud Compare 3D point cloud registration software. A standardized process was implemented to align the models to their bounding box centers and perform fine registration (*22*, *52*). A custom LUT centered around 0 was implemented to map positive (red) and negative (blue) deviations from the intended .STL onto the segmented .STL model. Gauging data and rendered 3D images of the deviations were exported for further analysis, display, and graphing.

Manual Perfusion of Branching Vascular Bed CHIPS

Perfusion of the branching vascular bed CHIPS was demonstrated by manually perfusing a concentrated dd$H_2O$ solution of blue food coloring through the top inlet of the construct with a 10 mL plastic BD syringe and a 20-gauge needle. The perfusion rate was modulated to achieve filling of the branching network and exit from the bottom outlet. Micromanipulators and helping hands devices were implemented to stabilize the perfusion without moving the printed construct. Perfusion was performed with the vascular bed construct submerged in a 50 mM HEPES buffer, pH 7.4.

VAPOR Assembly

Vasculature and perfusion organ-on-a-chip reactor (VAPOR) was designed with computer-aided design (CAD) software (Autodesk, Inventor; Autodesk, Fusion 360) and printed from Biomed Clear (Formlabs, RS-F2-BMCL-01) resin on a Form 3B (Formlabs, RS-F2-BMCL-01). Stainless steel M3 hex nuts (McMaster Carr, 94150A325) were inserted into the nut cutouts in the bioreactor main body. A glass coverslip was sealed into the lid by pipetting on 100 μL of resin followed by pressing a 22 x 22 mm glass coverslip (VWR, 48366-227) into the lid which was then sealed in place by baking in a UV oven. Custom gaskets were cut from 1.5 mm thick silicone sheeting (McMaster-Carr, 5787T93) and pressed into a cutout in the lid.

Bioreactor Perfusion System Assembly

A 100 mL glass bottle (Cole Parmer, EW-34523-00) was used as a media reservoir. Holes for tubing lines were drilled into the cap and 1/16" ID silicone tubing (Cole Parmer, EW-95802-02) was pulled through to ensure airtightness. Two additional holes were drilled for air filtration and media exchange. A peristaltic pump (Ismatec, EW-95663-34) using 1.42 mm ID peristaltic tubing (Cole Parmer, EW-95663-34) was connected to the media reservoir tubing. Autoclavable bubble traps (Darwin Microfluidics, LVF-KBT-L-A) with 1/16" barb adapters (Darwin Microfluidics, CIL-D-646) were placed after the pump and then connected to stopcocks on the reactor. Tubing exiting the reactor's perfusion channels and lymph outlet returned to the media reservoir.

Bioreactor Perfusion of CHIPS

For sterile operation 3D printed parts were sonicated for 30 min in sterile filtered 70% ethanol, dried for 1 hour in a biosafety cabinet, and sterilized by 15 min UV ozone treatment. All remaining parts such as peristaltic tubing, media reservoir and bubble traps were autoclaved. For air filtration, a 0.2 μm pore size filter (VWR, 28145-501) was screwed onto the media reservoir. The system was assembled in a biosafety cabinet on an incubator shelf. All flow paths were purged with media to avoid entrapment or perfusion of air bubbles on or in the tissue. CHIPS were then transferred into the VAPOR chamber, gently pressed onto the barbs and sealed by bolting on the lid. CHIPS were perfused from 60 – 1000 μL/min. The system was then inserted into a 37$^{o}$C incubator for perfusion culture. Cellularized CHIPS containing vascular bioink (HUVEC and MSC) were perfused with endothelial cell media while pancreatic CHIPS were perfused with a 50:50 ratio of endothelial cell and MIN6 media.(*53*)

pH-Sensitive Dye Perfusion

Serpentine CHIPS were placed in VAPOR reactors and perfused at 100 μL/min at each inlet. Colorimetric video acquisition was performed with a Sony A5 camera equipped with a Laowa 24mm f/14 probe lens. During perfusion, one flow path consisted of an acidic phenol red solution adjusted to pH 6.5 using hydrochloric acid (HCl). The second flow path consisted of a basic PBS buffer solution adjusted to pH 11 using sodium hydroxide (NaOH). Initial perfusion was performed with a pulsatile roller pump (Masterflex 77202-60) to stimulate mixing along the serpentine network length. The change in phenol red color from the acidic yellow to basic magenta was quantified using the Color Profiler (ImageJ (*53*), downloaded and installed from https://imagej.net/ij/plugins/color-profiler.html) for a segmented line then traversed the length of the serpentine network. Values for Magenta (White-Green) and Yellow (White-Blue) were extrapolated from RGB intensity to determine the ratio of Yellow:Magenta along the path during different perfusion states. The ratio was graphed as a function of path length along the serpentine network and color coded to match the yellow and magenta values according to the pH indicator values for phenol red.

Color Dye Perfusion

Laminar flow within Serpentine CHIPS was demonstrated by perfusing either red or blue food coloring while maintaining equal flow rates of 100 µL/min in both channels. Vessel patency and dye diffusion into the bulk of stacked or 3D helical channel CHIPS was demonstrated by perfusing red and blue food coloring through separate channels. Colorimetric video acquisition was performed with a Sony A5 camera equipped with a Laowa 24mm f/14 probe lens. Extended time lapse imaging was acquired with a GoPro Hero 5 camera mounted to a tripod.

FITC-Conjugated Dextran Perfusion

Dual parallel channel CHIPS were perfused at 100 µL/min with 0.1 mg/mL 3, 10, 40 or 70 kDa dextran. Dextrans were conjugated with fluorescein isothiocyanate (FITC) (Thermofisher Scientific, D3305; D1821; D1844; D1823). Dual parallel channel CHIPS' second vessel was perfused with 1X PBS. CHIPS were perfused from 1 – 72 hours. Time lapse images were recorded on an epifluorescent stereomicroscope (either Nikon SMZ1000; or Leica M165FC) using a FITC filter, an X-Cite lamp (Excelitas), and a Prime 95B Scientific CMOS camera (Photometrics) with an image being taken every minute. To quantify dextran diffusion, fluorescence intensity over time was measured using ImageJ (National Institutes of Health) (*53*). Six regions of interest (ROIs) were selected at increasing distances away from the FITC and PBS channels and fluorescence intensity over time was calculated relative to the intensity at the initiation of perfusion while accounting for background signal.

Microbead Perfusion

Fluorescent polystyrene microbeads 10 µm in diameter were perfused at 100 µL/min at a concentration of 3.6 X $10^3$ beads/mL. Beads had either 580/605 (red) (Thermofisher Scientific, F8838) or 505/515 (yellow-green) (Thermofisher Scientific, F8836) excitation/emission wavelengths. Beads were perfused through dual parallel CHIPS in either the same or opposite directions, and videos were recorded on stereofluorescence microscopes with a TexasRed filter set similar to dextran perfusions. Particle tracking and bead velocimetry were performed in Imaris 9.5.1 (Bitplane) using spot detection and tracking algorithms.

Perfusion of Dual Parallel Channel CHIPS with Afterload Pressure

Dual parallel channel CHIPS were perfused as previously described at 100 μL/min with 0.1 mg/mL 40 kDa FITC-conjugated dextran and 1X PBS. Each reservoir contained 20 mL of solution. Pressure within the CHIPS' dextran channel was increased by raising the height of the dextran reservoir to produce an additional 5 or 10 mmHg of afterload. To assess the diffusion of dextran from the source channel into the systemic PBS circulation, 50 μL samples were taken from the PBS reservoir bottle at 0 and 24 hours. The relative concentration of FITC-conjugated dextran compared to the source reservoir was then assessed by spectrophotometric analysis (Molecular Devices, SpectraMax i3x).

To assess the effect of afterload on molecular diffusion through CHIPS, time lapse images of perfusion with 5 mmHg of afterload (HP) were recorded as previously described and compared to perfusion with no additional afterload pressure (NP). The recordings were overlaid, and the fluorescence signal of the HP time lapse was divided by the NP time lapse after accounting for background signal. A vertical profile analysis was performed down the center of the HP/NP time lapse in ImageJ at various time points to further visualize the impact of increased afterload on diffusion into the peripheral regions of CHIPS.

Multi-Material Needle Alignment

In order to align multiple needles, we created a custom dual camera optical alignment system. Briefly, two 1X, 40mm WD CompactTL™ Telecentric C-mount Lens (Edmund Optics #63-745) were mounted to Alvium 1800 U-500 (Allied Vision) USB cameras. A custom 3D printed alignment plate and XY positioning system allowed for focus adjustment to achieve parfocality. To image the bottom needle tip to obtain the XY position and needle diameter, a mirror (Thorlabs ME2S-G01) was mounted at a 45° angle. The second camera was mounted perpendicular to the XY camera to view the side profile of the needle tip for Z-height alignment. A custom LabView program was written to simultaneously view the XY and Z positions. Each extruder was then moved to the center of the field of view for each camera to measure the relative XYZ offsets between each needle. The offset positions were stored as additional global software coordinate systems for use during multi-material printing using the Aerotech CNC operator's interface.

Multi-Material FRESH Printing

3D models were prepared using Fusion 360 (Autodesk) for multi-material printing by creating individual nested components for each material within the desired location of the CHIPS. Each component was exported as an STL part file and imported into Cura 5.2 (Ultimaker) slicing software. The main components were centered around the XYZ origin and offset according to their designed spacing based on the original 3D model location. A separate material profile was created for each bioink within Cura to permit the assignment and indexing of the respective bioinks to one of the three extruders. Additionally, the creation of individual bioink specific profiles enabled component-specific color visualization of the CHIPS within the slicing software. Custom start and end G-code was specified for each extruder tool profile to recall the stored position offsets determined during the alignment process and prime the extruder between tool changes.

Immunofluorescence Staining

Cellularized CHIPS were fixed via incubation in 10% neutral buffered formalin (Sigma Aldrich, HT501128) supplemented to a final molarity of 630 μM $MgCl_2$ and 108 μM $CaCl_2$. Tissues were then incubated in blocking buffer overnight. Blocking buffer consisted of 90% (v/v)

1X PBS supplemented to a final molarity of 1 mM $CaCl_2$ and $MgCl_2$ each, 5% (v/v) 1M Glycine (Fisher Scientific, BP381), 5% (v/v) goat serum (Thermo Fisher Scientific, 16210072), and 0.1% (v/v) Triton X-100 (Thermo Fisher Scientific, 85112). Samples were then immediately incubated with primary antibodies for a week at 4°C. Primary antibodies are diluted in antibody dilution buffer consisting of 1X PBS supplemented to a final molarity of 1 mM $CaCl_2$ and $MgCl_2$ each, 0.1% (w/v) bovine serum albumin (BSA) (Sigma Aldrich, A2153), and 0.1% (v/v) Triton X-100. Primary antibodies and their dilutions included VE-Cadherin rabbit mAb (Cell Signaling Technology, 2500S) at 1:400, CD-31 mouse mAb (Cell Signaling Technology, 3528S) at 1:800, insulin mouse mAb (Cell Signaling Technology, 8138S) at 1:400, and insulin rabbit polyclonal (Abcam, ab181547) at 1:400. Samples were then washed 3 times for 1 hour in antibody buffer without triton followed by an overnight wash at 4°C. The following day, samples were incubated in secondary antibodies for a week at 4°C. All secondary antibodies were diluted in antibody dilution buffer. Secondary antibodies and their dilutions included 4′,6-diamidino-2-phenylindole (DAPI) (Sigma Aldrich, D9542) at 1:400, phalloidin conjugated to Alexa-Fluor 488 (Life Technologies, A12379) at 1:400, goat anti-rabbit IgG 555 (Thermofisher Scientific, A- 21428) at 1:1000 dilution, and goat anti-mouse IgG 633 (Thermofisher Scientific, A-21050) at 1:1000. Samples were then washed 3 times for 1 hour in antibody buffer without triton followed by an overnight wash at 4°C.

Tissue Clearing

After immunofluorescent staining, tissues were optically cleared using Benzyl Alcohol/ Benzyl Benzoate (BABB). Samples were first serially dehydrated by 1 hour incubation each in 10%, 25%, 50%, 75%, 90%, and 100% (v/v) ethanol solutions. Samples were then transferred into fresh 100% ethanol solution and incubated overnight at 4°C. Samples were then optically cleared by incubation in BABB for at least 1 hour prior to imaging.

Confocal Imaging

All fluorescence confocal imaging was performed on a Nikon A1R HD MP multiphoton microscope equipped with a 4× (NA = 0.20) plan apochromat objective, 16× (NA = 0.80) long working distance water immersion objective, a 25× (NA = 1.10) plan apochromat water immersion objective, 4 visible light internal detectors, 4 visible laser lines (405, 488, 561, 633 nm), a motorized Prior Z-deck stage, Piezo Z Nosepiece, and Insight X3 DeepSee multiphoton laser (Spectra Physics). Large overview tile scans and 3D z-stack images of tissues were acquired using the 4× (NA = 0.20) plan apochromat (Nikon) objective with NIS Elements software. 3D rendering and image processing was performed in Imaris (v9.5, Bitplane). For cleared tissues, a custom machined aluminum chamber was constructed to permit imaging through a large coverslip window and immobilization of the tissue within the BABB solution.

Fluorescence Image Analysis

Advanced 3D fluorescence image analysis and animations were generated in Imaris 10.0 (Oxford Instruments). Specifically, the machine learning based vascular segmentation wizard was utilized to quantify the migratory network density and diameter within the vascular CHIPS. Colocalization analysis for the Actin channel with CD-31 channel in the pancreatic CHIPS was performed on regions of interest using the Coloc-2 plugin within FIJI Image-J. Pearson's correlation coefficients and manders M1 and M2 values were recorded.

Glucose-Stimulated Insulin Secretion

Glucose stimulated insulin secretion (GSIS) was conducted using a static incubation approach in which FRESH bioprinted pancreatic CHIPS were removed from the bioreactor. The MIN6-containing pancreatic CHIPS were exposed to serial incubations in 4 mL low glucose (LG) (1.67 mM) and high glucose (HG) (16.7 mM) Krebs Buffer similar to previous work (*36*, *54*, *55*). A pre-incubation period in LG Krebs buffer was followed by serial incubations in fresh LG followed by HG Krebs buffer for 1.5 hours each. GSIS was performed in duplicate. Samples collected from each incubation phase were stored at -80°C for subsequent analysis of insulin concentration. Insulin content was analyzed using a mouse insulin ELISA (Mercodia) with each sample assayed in duplicate.

Statistics and Data Analysis

Statistical analysis was performed with Prism 10 (Graphpad) using appropriate tests based on experimental conditions and data. For comparison of GSIS insulin concentrations, an unpaired t-test was performed. Statistical significance was based on a $P < 0.05$ (*) with lower P-values being denoted as $P < 0.01$ (**). Non-significant P-values were denoted as ns.

Figures and visuals were constructed using Illustrator version 28.1 and Photoshop version 25.3.1 (Adobe). Supplemental videos and time-lapse images were edited in FIJI ImageJ and compiled in Premiere Pro version 24.1 (Adobe).

Supplementary Figures:

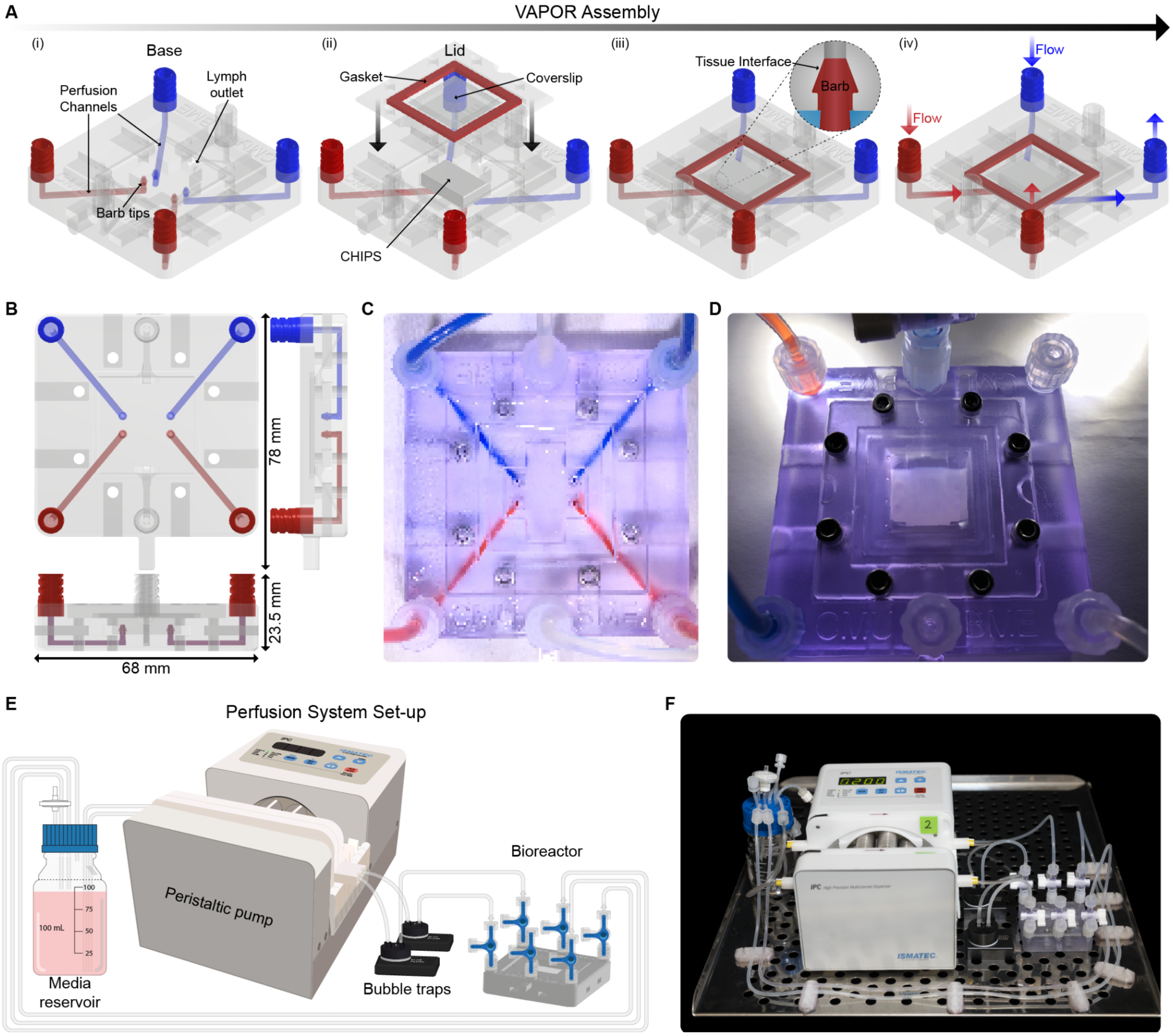


**fig. S1. Design and implementation of the VAPOR bioreactor platform.** (**A**) CAD model and schematic of the (i) VAPOR base, (ii) lid and gasket assembly, (iii) CHIPS and VAPOR barb interface for watertight seal, (iv) dual independent flow paths through VAPOR and CHIPS. (**B**) Dimensions of the assembled VAPOR system. (**C**) Image of VAPOR perfused with red and blue dye to visualize the internal fluidic networks. (**D**) Fully assembled VAPOR and inserted serpentine CHIPS prior to perfusion. (**E**) Schematic of the perfusion setup utilized in VAPOR. (**F**) Image of the assembled VAPOR platform and perfusion system prior to installation into a cell culture incubator.

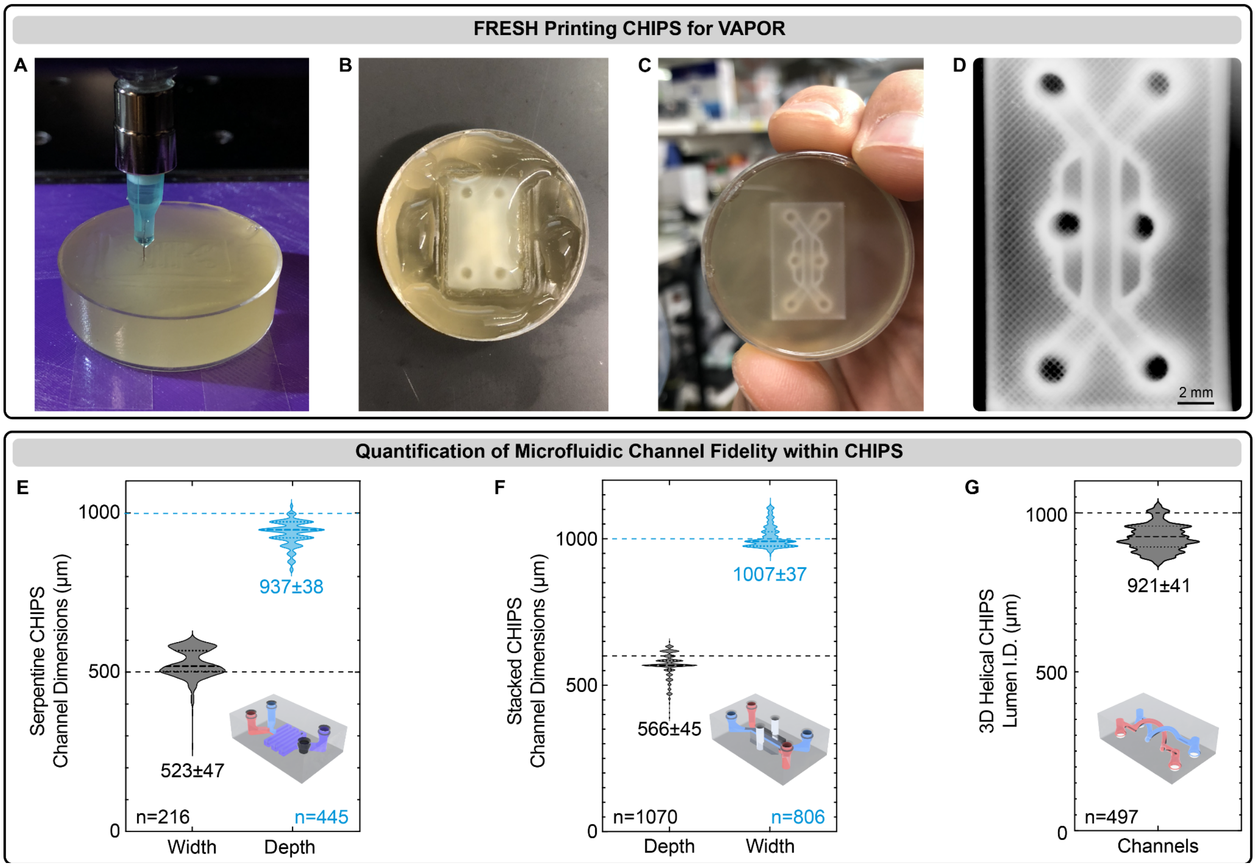


**fig. S2. FRESH printing CHIPS designed for VAPOR perfusion and internal network quantification.** (**A**) A CHIPS model midway through FRESH printing within a gelatin support bath. (**B** and **C**) Top (B, helical) and bottom (C, stacked) view of CHIPS immediately after FRESH printing. (**D**) A stereomicroscope image of a Stacked CHIPS model after release from the FRESH support bath. (**E** to **G**) OCT quantification of measured versus expected channel dimensions for Serpentine (G), Stacked (H), and 3D Helical (I) CHIPS.

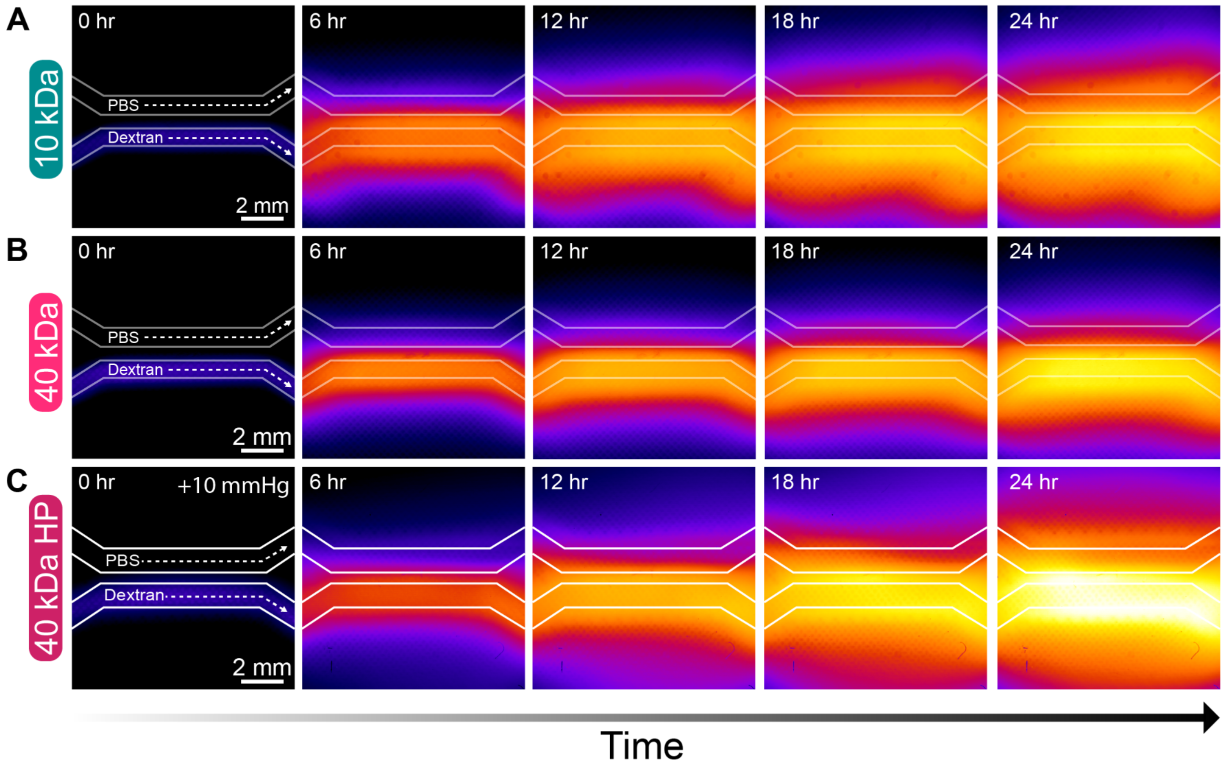


**fig. S3. Additional diffusivity analysis within dual channel CHIPS.** (**A**) Time lapse images of dual parallel channel CHIPS undergoing VAPOR perfusion with FITC-conjugated 10 kDa dextran. (**B**) Time lapse images of dual parallel channel CHIPS undergoing VAPOR perfusion with FITC-conjugated 40 kDa dextran under normal pressure. (**C**) Time lapse images of dual parallel channel CHIPS undergoing high pressure (+10 mmHg) VAPOR perfusion of 40 kDa FITC-conjugated dextran over 24 hours.

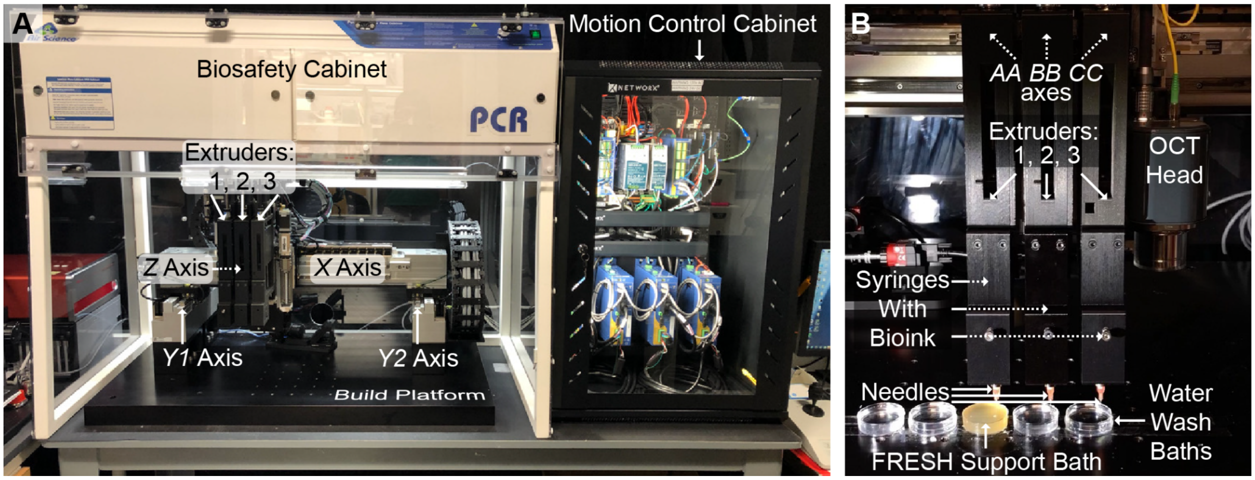


**fig. S4. Design and implementation of a multi-material bioprinter.** (**A**) Custom high-performance 3D bioprinter based on a commercial Aerotech motion control platform and open-source Replistruder 5 syringe pumps. (**B**) Three Replistruder 5 syringe pumps are utilized for material multi-material FRESH printing of CHIPS. An onboard OCT system allows for volumetric imaging and in-process monitoring of FRESH printed CHIPS.

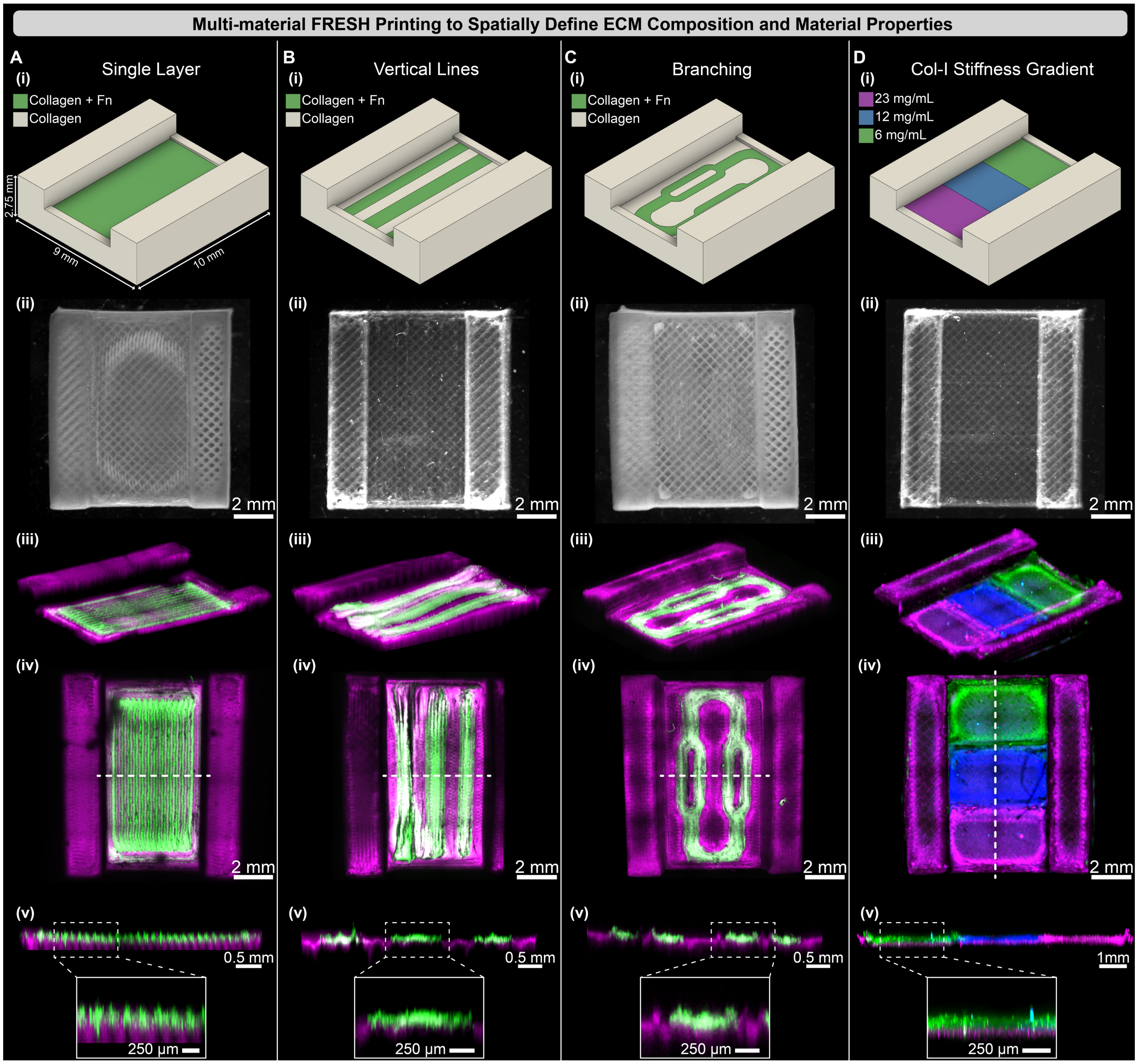


**fig. S5. Multi-material FRESH printing to volumetrically define ECM composition, material properties, and growth factor localization.** (**A** to **C**) A parallel plate style CHIPS design with a collagen frame and uniform single layer (A), vertical lines (B), and branching network (C) of fluorescently-tagged fibronectin (Fn) containing collagen displayed as (i) CAD design and stereomicroscopy image, and (ii) quantified for multi-material printing registration with 3D confocal microscopy (iii-v, dotted line indicates cut plane, inset scale bar in v = 250 µm). (**D**) A collagen stiffness gradient generated by multi-material printing of 6, 12, and 23 mg/mL collagen in adjacent regions within the parallel plate CHIPS (i), stereomicroscopy image (ii) of the 3-material print demonstrates high-fidelity and multi-material registration upon fluorescence image analysis (iii-v).

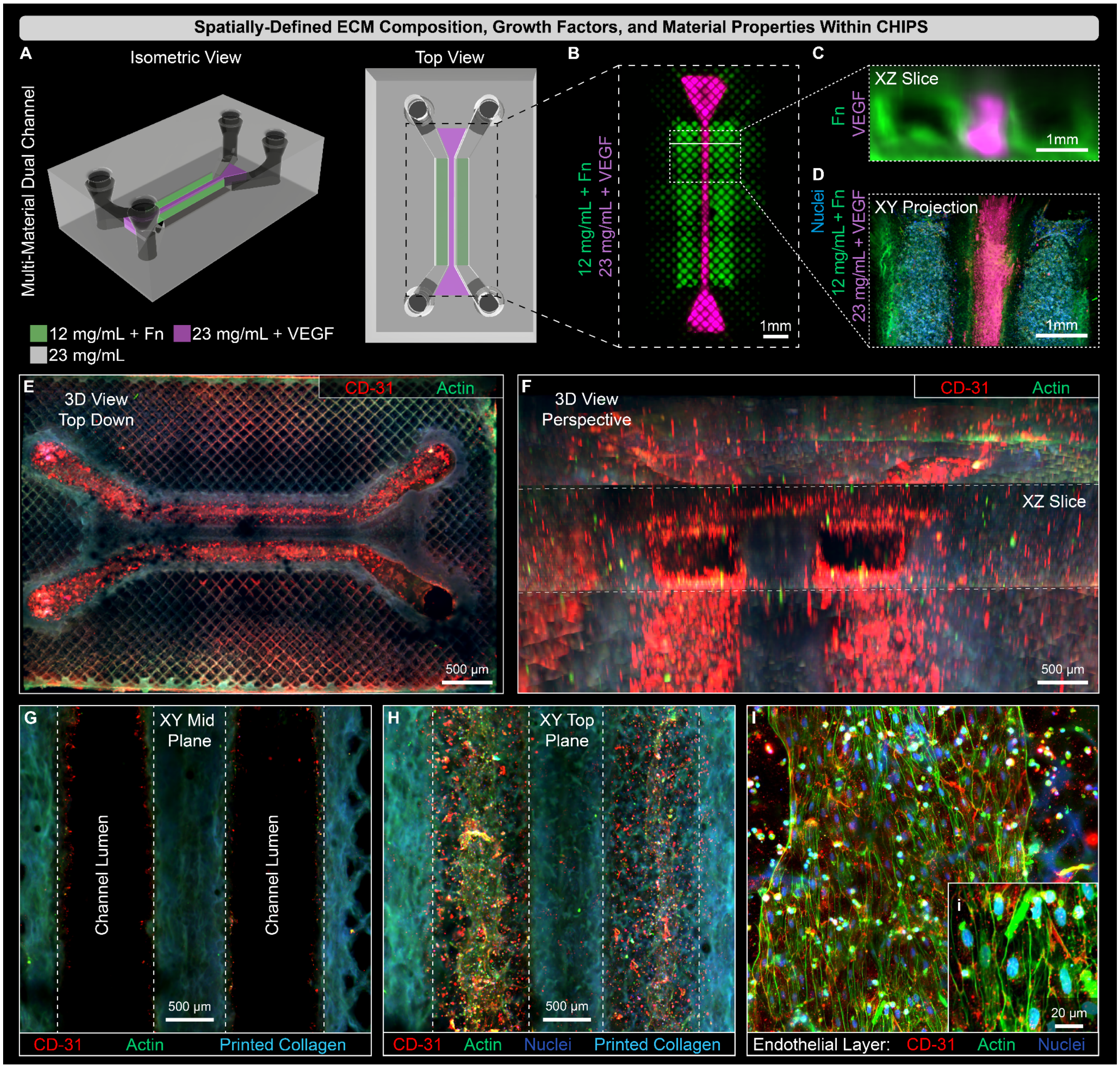


**fig. S6. Perfusion seeded endothelial cells form monolayer within multi-material CHIPS.** (**A**) Dual channel CHIPS CAD design of internally lined channels containing 12 mg/mL collagen + Fn (Green) with a central dividing region between the fluidic channels containing 23 mg/mL collagen + VEGF growth factor (Magenta). (**B**) Confocal fluorescence maximum Z-projection image of the 12 mg/mL collagen channel lining and VEGF patterning. (**C**) XZ slice plane image from (B) demonstrating the Fn channel lining (green) and VEGF patterning (Magenta). (**D**) Confocal fluorescence image of central channel regions from (B) seeded with HUVECs and stained for nuclei distribution (blue) at day 1 after seeding. (**E**) Volumetric confocal fluorescence imaging of optically cleared multi-material dual parallel channel CHIPS stained for endothelial cell marker CD-31 (Red) and actin (Green). (**F**) A 3D view with XZ cross-section of endothelial channel lining (Red). (**G**) XY mid-plane view of open channel lumen and cell seeding. (**H**) XY top surface of channel view of endothelial monolayer coating the channel lumen. (**I**) Magnified XY images of endothelial lining within channels and increased zoom (i).

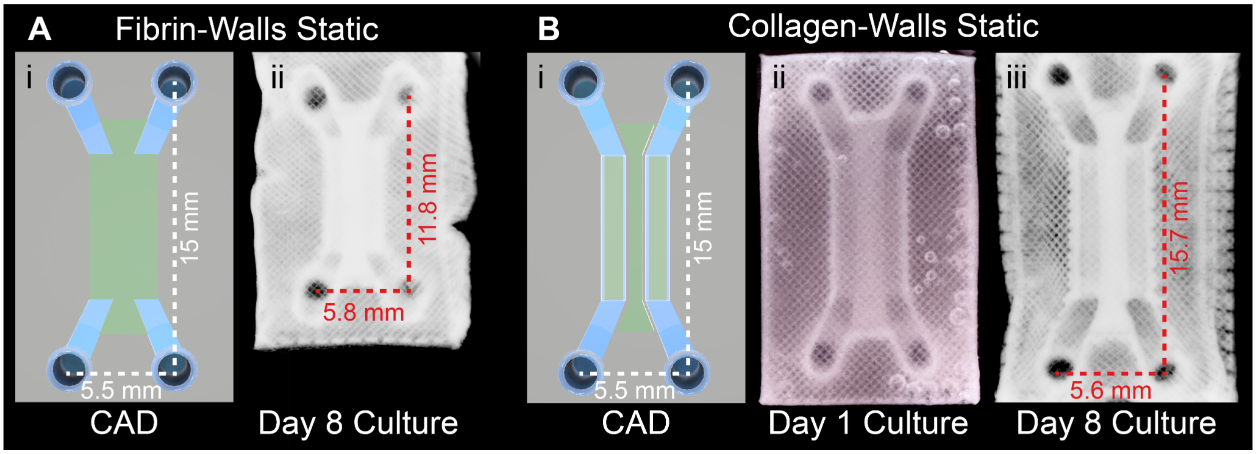


**fig. S7. The addition of collagen walls lining the perfusion channels prevent cell-driven buckling.** (**A**) Dimensional comparison of the vascular CHIPS with complete cellular + fibrin walls CAD model (i) to the vascular CHIPS after 8-days of static culture (ii). (**B**) Dimensional comparison of the vascular CHIPS with collagen walls CAD model (i) to vascular CHIPS after 1 (ii) and 8-days of static culture (iii).

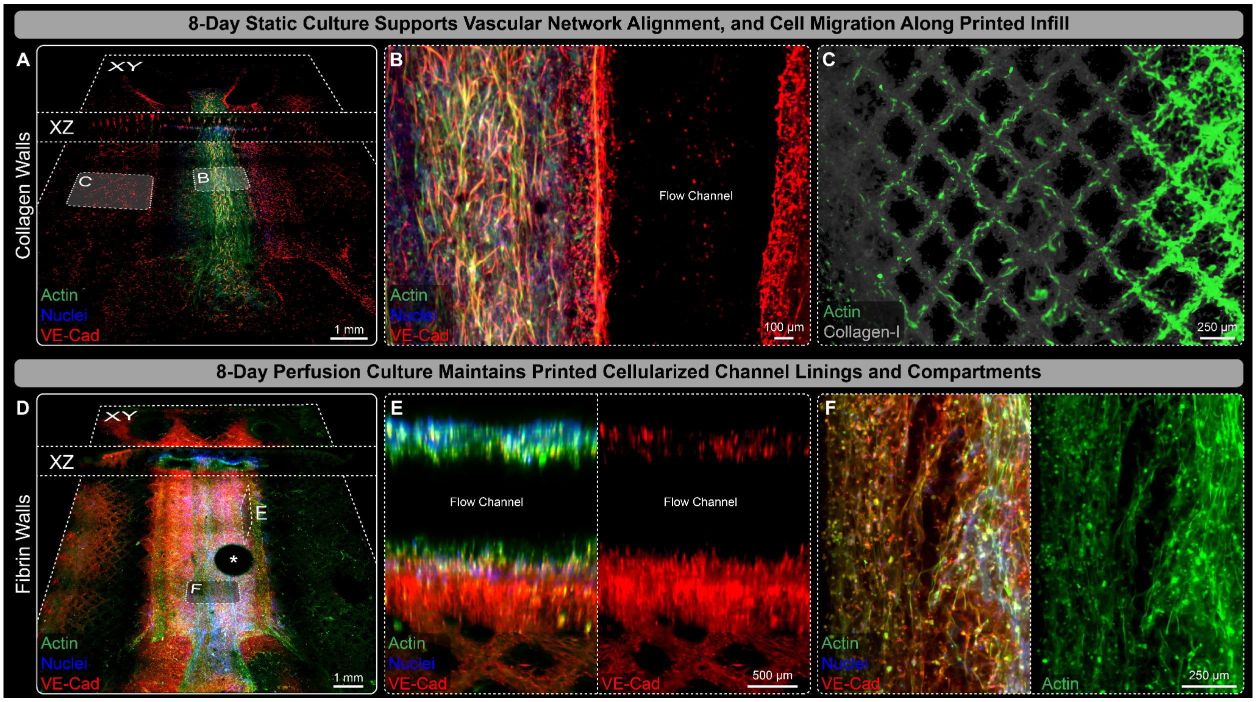


**fig. S8. FRESH printed cellularized channel linings and directed cell migration along printed collagen filaments within CHIPS.** (**A**) Both XY and XZ perspective slice views of the channel bottom surface from 3D confocal imaging of optically cleared dual parallel channel cellular CHIPS with collagen walls statically cultured for 8 days. (**B** and **C**) Images reveal cellular alignment along the length of the channels, lumenal VE-Cadherin expression (B), and evidence of cell migration following the printed collagen infill of distances exceeding 1.5 mm from the channel outside edge (C). (**D**) Both XY and XZ perspective midplane slice views from 3D confocal imaging of optically cleared dual parallel channel fibrin walls cellular CHIPS following 8 days of perfusion culture within VAPOR. (**E** and **F**) Images reveal enhanced VE-Cadherin expression around the flow channels (E), and extensive cell spreading throughout the central cellular region between channels (F). *Indicates air bubble artifacts introduced into the channels during CHIPS optical clearing and imaging.

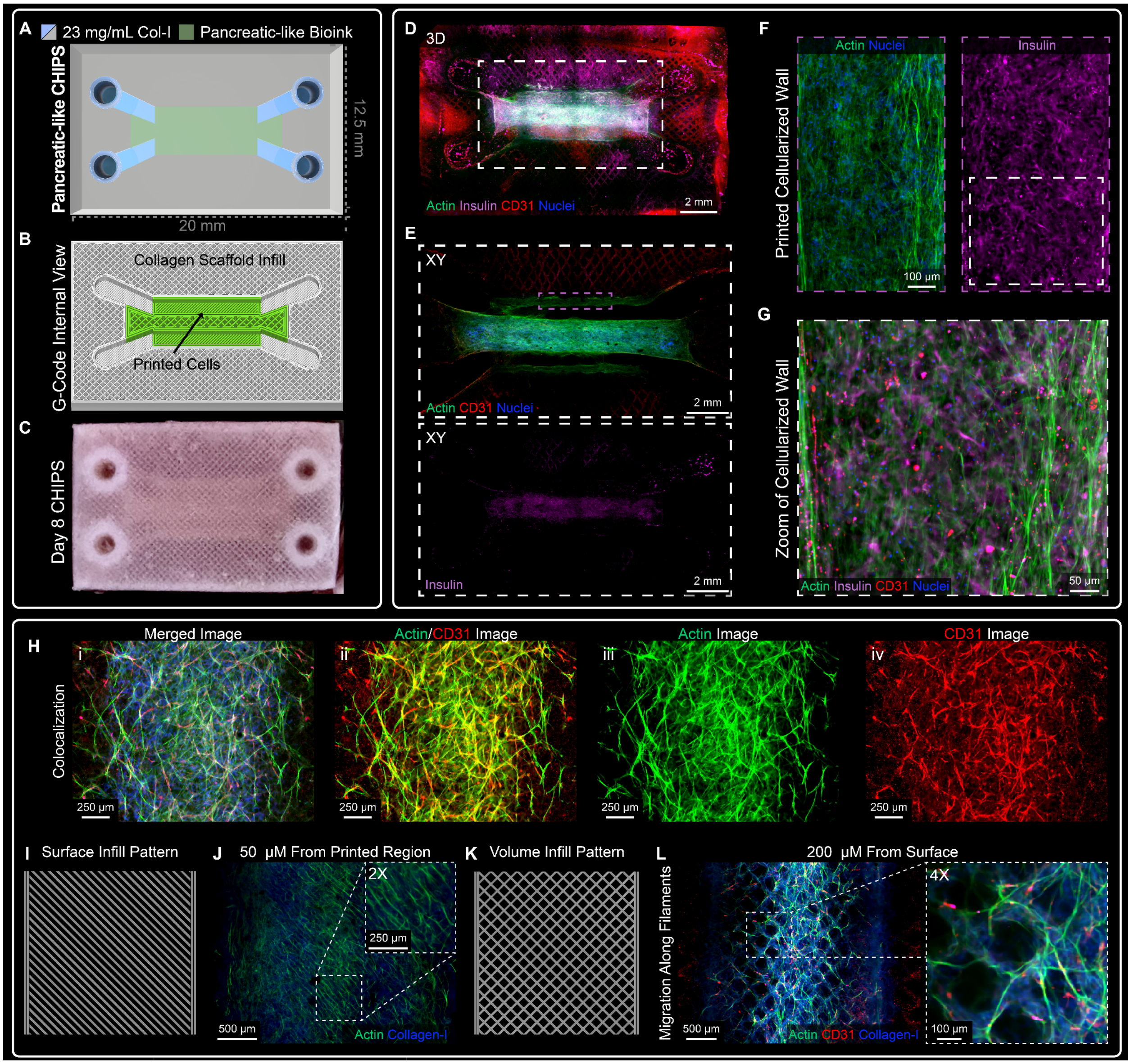


**fig. S9. Pancreatic CHIPS with fibrin walls express insulin and show a high degree of vascular migration along collagen filaments. (A** and **B)** Schematic design and machine pathing G-code of pancreatic CHIPS with fibrin walls. **(C)** Stereomicroscope image of a pancreatic CHIPS after 8 days of static culture. **(D)** XY plane view from whole mount confocal fluorescence imaging of optically cleared 8-day statically cultured pancreatic CHIPS. **(E)** Zoomed region from (D) of the central cellularized area within the CHIPS. **(F)** Zoomed region of the printed channel wall from (E) highlighting the high cellularization achieved via 60 million cell/mL bioink and presence of insulin expression. **(G)** Zoom region from (F) showing the cellular distribution and protein expression within the fibrin printed wall after 8 days of static culture. **(H)** Confocal fluorescence imaging highlighting CD-31 colocalization with Actin within CHIPS resulting in a Pearson's Correlation Coefficient of $0.63 \pm 0.04$. **(I)** Graphic illustration of the FRESH printing pattern at the surface of the cellular region. (**J**) Confocal fluorescence image of the cellular alignment (actin, green) along the direction of the original print path (-45° angle) shown in (I). (**K**) Graphic illustration of the FRESH printed volume infill pattern (alternating ±45° angle) at a depth 200 µm from the surface of the cellular region. (**L**) Cellular migration (actin, green) along the printed collagen filaments (blue) 200 µm below the original printed cellular region.

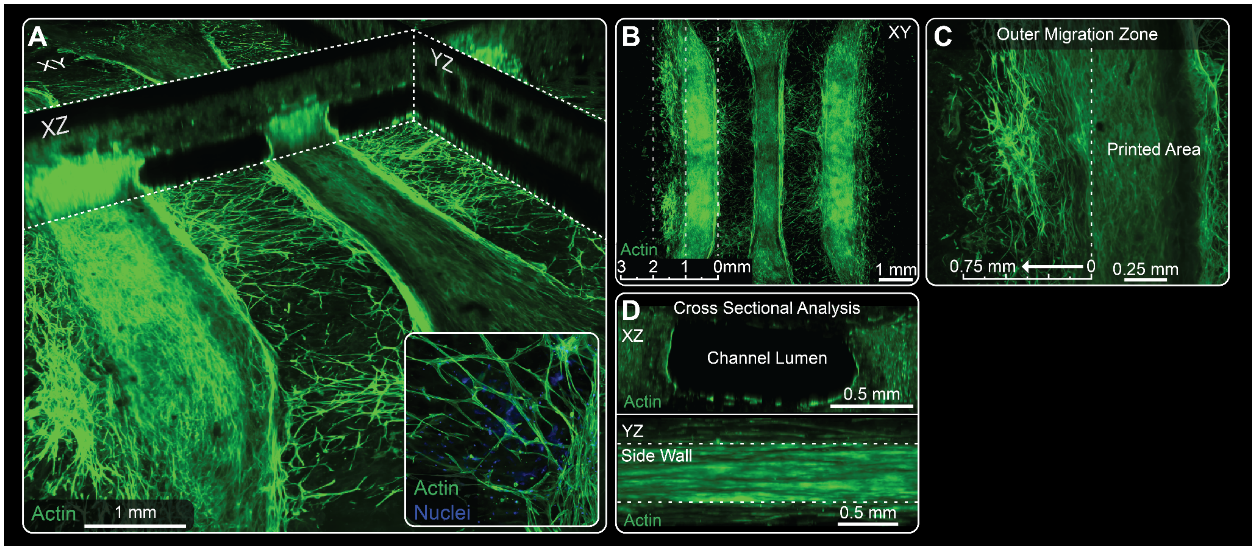


**fig. S10. Highly migratory and branching cellular networks within statically cultured pancreatic CHIPS.** (**A**) Confocal fluorescence images of XY, YZ, and XZ plane views revealing the intricate cell network and migration between the patent flow channels. (**B**) XY max intensity Z-projection image demonstrating the range of cell migration (actin, green) outward from the printed regions. (**C**) Zoom in image to the outer migration zone. (**D**) XZ cross sectional analysis expression profile of the cellular (actin, green) markers lining the original acellular channels, in addition to a YZ projection showing a dense luminal cell monolayer along the side walls.

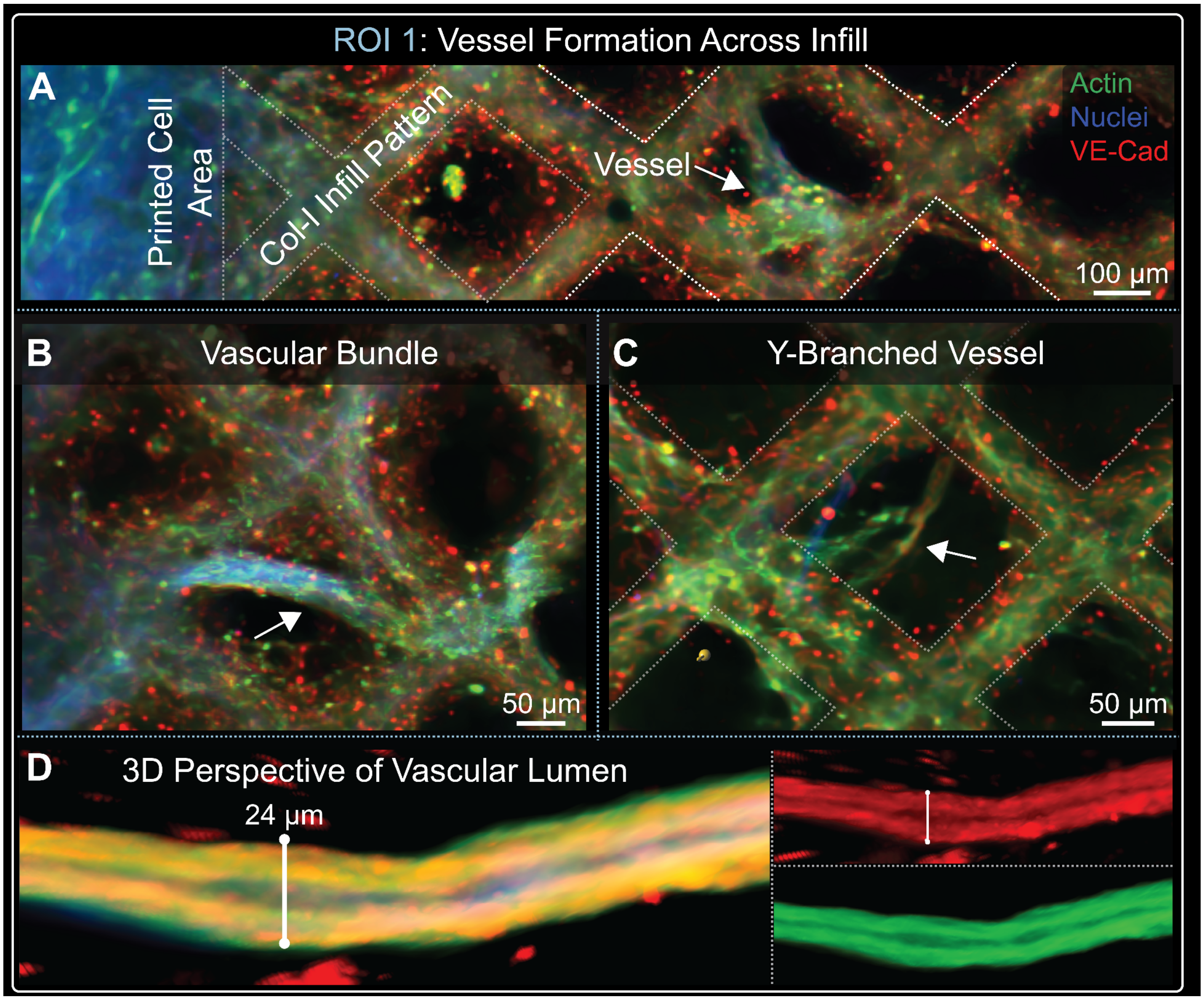


**fig. S11: Vasculogenesis and formation of branching networks across printed infill patterns.** (**A** to **C**) Confocal fluorescence images from ROI 1 in Figure 5 (E) demonstrating a (A) branched vessel-like structure 100 µm in diameter, (B) A vascular-like bundle of 50 µm in diameter bridging and following the infill lattice structure, and (C) a 25 µm Y-branched vascular-like structure bridging across the collagen infill pattern (outline with dotted white lines) within the CHIPS. (**D**) 3D perspective view of a 24 µm vessel with visible open lumen expressing actin (green) and VE-cadherin (VE-Cad, red).
